# Morphologic intratumoral heterogeneity from routine whole-slide histopathology is prognostic for survival in primary central nervous system lymphoma: development in the LOC Network and international external validation

**DOI:** 10.64898/2026.08.31.748241

**Authors:** Lucas Rincón de la Rosa, Noémie Barillot, Isaias Hernández-Verdin, Roser Velasco, Bertrand Mathon, Sandrine Eimer, Jean Rodolphe Vignes, Audrey Rousseau, Jérôme Paillassa, Guido Ahle, Felix Lerintiu, Emmanuelle Uro-Coste, Lucie Oberic, Emeline Tabouret, Romain Appay, Guillaume Gauchotte, Luc Taillandier, Jean-Pierre Marolleau, Clovis Adam, Renata Ursu, Stefania Cuzzubbo, Anna Schmitt, Lucia Nichelli, Albert Pons-Escoda, Frédéric Charlotte, Frédéric Davi, Magali Le Garff-Tavernier, Sylvain Choquet, Carole Soussain, Noemí Vidal, Eva González-Barca, Fina Climent, Patricia López, Fanny Drieux, Elena-Liana Veresezan, Michael Heming, Gerd Meyer zu Hörste, Oliver Grauer, Fabrice Jardin, Karima Mokhtari, Caroline Houillier, Khê Hoang-Xuan, Agustí Alentorn

## Abstract

**Background:** Clinical scores incompletely capture outcomes in primary central nervous system lymphoma (PCNSL). We quantified morphologic heterogeneity in pretreatment hematoxylin and eosin (H&E) whole slides.

**Patients and methods:** Three independent cohorts of immunocompetent, HIV- and EBV-negative patients treated recently were analyzed: LOC 2023 (122 slides), phase III BLOCAGE-01 (245 slides; NCT02313389), and external Barcelona (BCN; 41 slides). UNI embeddings, prototype learning, spatial metrics, and elastic-net Cox regression defined ITH-C.

**Results:** Models achieved bootstrap-corrected concordance of 0.797–0.834. Age-, sex-, and KPS-adjusted ITH-C HRs were 1.29 (95% CI 1.01–1.64), 1.27 (1.07–1.51), and 2.13 (1.35–3.37), respectively. Adding ITH-C increased MSKCC C-index from 0.671 to 0.717, 0.560 to 0.593, and 0.588 to 0.706. Spatial transcriptomics linked ITH-C to immune programs.

**Conclusions:** Routine H&E encodes prognostic spatial heterogeneity in PCNSL. ITH-C complements clinical scores, supporting prospective risk stratification.

## 1 Introduction

Primary central nervous system lymphoma (PCNSL) is a rare, aggressive extranodal non-Hodgkin lymphoma involving the brain, leptomeninges, eyes, or spinal cord. Most tumors are diffuse large B-cell lymphomas (DLBCLs) with activated B-cell or MYD88/CD79B-altered biology [1]. Outcomes remain heterogeneous despite high-dose methotrexate-based therapy and contemporary consolidation or maintenance approaches [2, 3]. The International Extranodal Lymphoma Study Group (IELSG) and Memorial Sloan Kettering Cancer Center (MSKCC) prognostic scores use clinical and laboratory variables [4, 5], but neither incorporates tissue morphology or spatial intratumoral heterogeneity (ITH). This omission is particularly relevant in PCNSL because diagnosis usually relies on a small stereotactic biopsy rather than a resection. The amount of available tissue is limited, repeated sampling is rarely justified, and conventional bulk assays reduce all cellular and spatial variation within the biopsy to one averaged measurement.

ITH encompasses coexisting genetic, transcriptional, epigenetic, phenotypic, immune, and spatial states. These forms of heterogeneity are related but not interchangeable. Genomic subclones can differ in treatment sensitivity; transcriptional states can change without fixed genetic divergence; and spatially restricted immune or stromal niches can alter the behavior of otherwise similar malignant cells [6, 7]. In TRACERx lung cancer, subclonal architecture and the evolution of copy-number alterations were associated with recurrence and outcome [8]. Multi-region studies have similarly connected evolutionary heterogeneity to prognosis in renal, ovarian, breast, head-and-neck, and brain tumors [9–11]. Pan-cancer analyses further suggest that the relationship is not necessarily linear: very low heterogeneity may reflect a highly fit dominant clone, whereas high heterogeneity may provide a broader repertoire for adaptation [12].

The tumor microenvironment introduces an additional layer. Immune-cell density alone does not describe whether immune populations are intermingled with malignant cells, excluded at a boundary, or confined to discrete vascular and stromal niches. Spatial organization may therefore contain clinically relevant information that is lost by bulk enumeration. Work on the immune contexture and spatially resolved immune escape has emphasized the prognostic importance of both composition and geography [13, 14]. In PCNSL, where lymphoma cells coexist with resident glia, recruited myeloid cells, reactive astrocytes, endothelial structures, and perivascular lymphocytes, a histology-derived heterogeneity measure should be interpreted as a tissue-ecosystem readout rather than as a tumor-cell purity measure. Histology offers a practical way to sample spatial phenotype at high resolution from routinely acquired tissue. Earlier computational pathology studies showed that stromal organization, cellular morphology, and local cellular neighborhoods in breast cancer were associated with outcome [15, 16]. Longitudinal risk was also linked to spatial heterogeneity of estrogen-receptor expression [17]. More recently, large-scale single-cell morphologic and topologic phenotyping characterized ecosystem diversity in breast cancer [18], while visual ITH features have been associated with tumor progression [19]. Deep-ITH demonstrated that transcriptional ITH inferred from routine breast histology retained independent prognostic information [20]; in glioblastoma, the GBM360 framework linked spatial cellular architecture to prognosis [21]. Together, these data support morphologic heterogeneity as a biologically meaningful intermediate phenotype rather than a purely visual descriptor.

DLBCL is itself heterogeneous across malignant B-cell states and microenvironmental niches. DLBCL-Morph showed that quantitative nuclear geometry from digital slides could support outcome modeling [22]. Multimodal spatial profiling of 78 DLBCLs identified seven recurrent cellular neighborhoods and substantial variation among and within tumors [23]; spatial transcriptomics has also resolved heterogeneous immune niches in EBV-positive DLBCL [24]. These findings are important for PCNSL because an anatomically restricted lymphoma can still contain several malignant and nonmalignant programs within one diagnostic specimen. In PCNSL, bulk transcriptomics defined four consensus signatures (CS1–CS4) that incorporate both tumor-cell and microenvironmental programs [25]. Single-cell and spatial transcriptomic profiling has independently demonstrated transcriptionally heterogeneous malignant B-cell clusters, spatially restricted enrichment, and colocalized T-cell exhaustion in PCNSL [26]. A bulk label identifies the dominant aggregate expression pattern, but does not establish that all regions of the biopsy share that state.

Computational pathology has moved from task-specific convolutional models toward transferable foundation representations. Weakly supervised models can learn from slide-level labels without exhaustive region annotation [27–29]. Self-supervised pathology encoders, including Phikon, UNI, Virchow, and Prov-GigaPath, capture reusable morphologic features across organs, stains, and downstream tasks [30–33]. H&E models can classify histologic entities, infer alterations such as driver mutations and microsatellite instability [34, 35], and predict survival across cancers [36, 37]. Prototype-based learning adds an interpretable intermediate representation by relating each patch to learned reference vectors, although prototype stability and biological identifiability remain important concerns [38].

We used a common WSI pipeline to quantify complementary aspects of morphologic ITH: prototype composition, confidence, local mixing, spatial autocorrelation, connected-region structure, and persistence across spatial scales. An elastic-net Cox combination was developed in pooled LOC 2023 and BLOCAGE-01 data, with signed coefficients retained for interpretation, and was then evaluated in BCN as an external international validation cohort. Spatial transcriptomics supplied an orthogonal RNA-based tissue readout for the H&E-defined phenotype.

## 2 Results

Figure 1 summarizes the computational pipeline. Whole-slide images and, for LOC 2023, bulk-RNA-seq-derived molecular subgroup labels were processed through a common CLAM tissue-segmentation and UNI patch-embedding trunk. A prototype-learning multiple-instance-learning model (TPMIL) was trained to predict the four bulk-derived consensus molecular subgroups while producing, for every tissue patch, a soft assignment to one of four tumor prototypes or a negative/background prototype. Applying the trained model yielded patch-level prototype maps that were summarized into slide-level composition and spatial-organization features. The same per-patch output, combined with the multiscale TP-HPM summary and elastic-net feature combination described in Methods, underlies the ITH-C composite. A companion survival model shared the preprocessing and UNI-embedding trunk but was trained directly for continuous risk prediction and evaluated by patient-level bootstrap resampling.

**Fig. 1.**
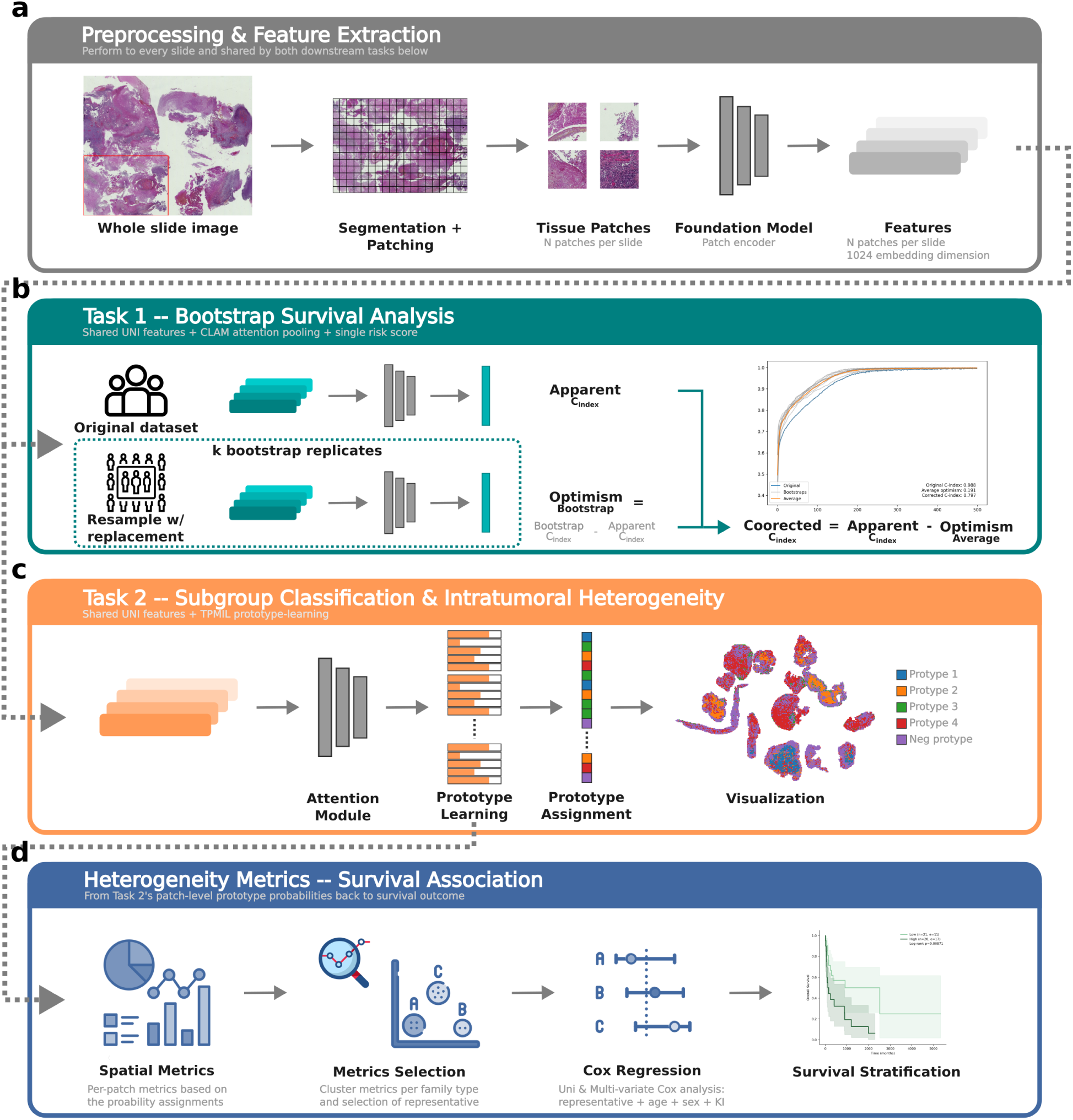
Whole-slide image processing and analytical workflow. **a)** Treatment-naive diagnostic H&E WSIs undergo CLAM tissue segmentation and patch extraction; UNI encodes each patch into a 1,024-dimensional vector. **b)** An attention-based Cox MIL model learns a continuous slide-level risk score separately within each cohort, with patient-level bootstrap optimism correction. **c)** TPMIL uses the same embeddings and LOC 2023 CS1–CS4 labels to learn four positive prototypes and one negative/background prototype; soft prototype probabilities are projected back to patch coordinates. **d)** Composition, diversity, neighborhood, spatial-autocorrelation, connected-region, and multiscale-persistence features are combined into ITH-C and evaluated with continuous Cox models.

### 2.1 Cohorts and analytic denominators

The source cohorts comprised 107, 244, and 40 patients in LOC 2023, BLOCAGE-01, and BCN, respectively. 2023 and BLOCAGE are often used in the figures legends instead of the complete cohort name. Some patients contributed more than one diagnostic slide, producing 122, 245, and 41 evaluable WSIs for slide-level analyses (Figure 2; Table 1). We retain both denominators throughout to distinguish clinical enrollment from image-level evaluation. All included patients were immunocompetent and HIV negative, and all PCNSLs were EBV negative by immunohistochemistry. BLOCAGE-01 was a multicenter randomized phase III study of maintenance rituximab–methotrexate–temozolomide versus observation in older patients with PCNSL who had achieved complete response after high-dose methotrexate-based induction (NCT02313389).

**Fig. 2.**
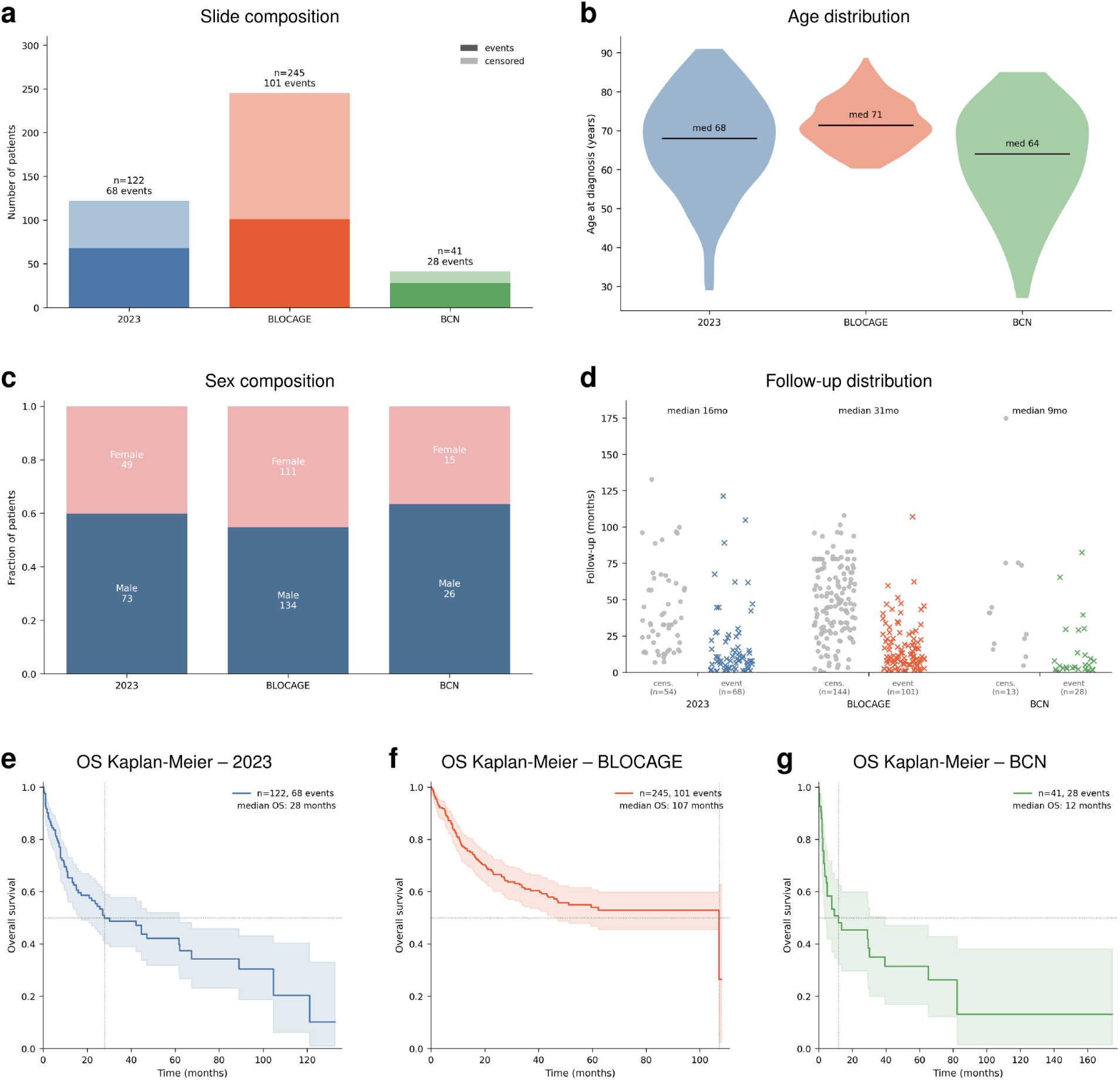
Cohort composition, demographics, and unadjusted overall survival. **a)** WSI-level events and censoring; totals denote evaluable slides. **b)** Age-at-diagnosis distributions; horizontal bars and annotations indicate median age. **c)** Sex composition. **d)** Observed follow-up by vital-status endpoint, with median observed time reported above each cohort. **e–g)** Kaplan–Meier OS curves with 95% confidence bands; median OS is displayed directly on each panel for LOC 2023 (**e**), BLOCAGE-01 (**f**), and BCN (**g**).

**Table 1:**
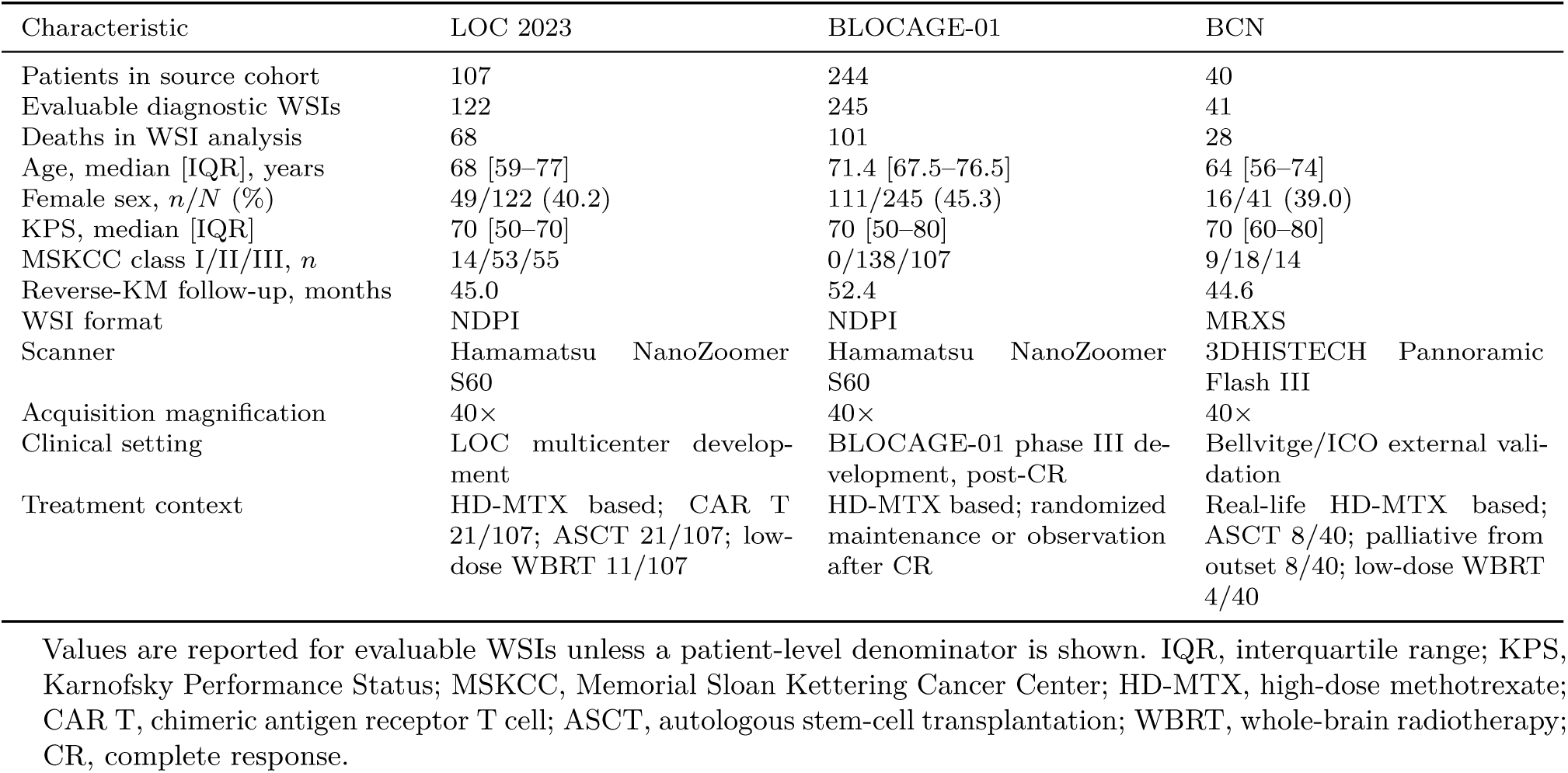
Baseline clinical, treatment, and image characteristics.

| Characteristic | LOC 2023 | BLOCAGE-01 | BCN |
| --- | --- | --- | --- |
| Patients in source cohort | 107 | 244 | 40 |
| Evaluable diagnostic WSIs | 122 | 245 | 41 |
| Deaths in WSI analysis | 68 | 101 | 28 |
| Age, median [IQR], years | 68 [59–77] | 71.4 [67.5–76.5] | 64 [56–74] |
| Female sex, $n/N$ (%) | 49/122 (40.2) | 111/245 (45.3) | 16/41 (39.0) |
| KPS, median [IQR] | 70 [50–70] | 70 [50–80] | 70 [60–80] |
| MSKCC class I/II/III, $n$ | 14/53/55 | 0/138/107 | 9/18/14 |
| Reverse-KM follow-up, months | 45.0 | 52.4 | 44.6 |
| WSI format | NDPI | NDPI | MRXS |
| Scanner | Hamamatsu NanoZoomer S60 | Hamamatsu NanoZoomer S60 | 3DHISTECH Pannoramic Flash III |
| Acquisition magnification | 40× | 40× | 40× |
| Clinical setting | LOC multicenter development | BLOCAGE-01 phase III development, post-CR | Bellvitge/ICO external validation |
| Treatment context | HD-MTX based; CAR T 21/107; ASCT 21/107; low-dose WBRT 11/107 | HD-MTX based; randomized maintenance or observation after CR | Real-life HD-MTX based; ASCT 8/40; palliative from outset 8/40; low-dose WBRT 4/40 |

Values are reported for evaluable WSIs unless a patient-level denominator is shown. IQR, interquartile range; KPS, Karnofsky Performance Status; MSKCC, Memorial Sloan Kettering Cancer Center; HD-MTX, high-dose methotrexate; CAR T, chimeric antigen receptor T cell; ASCT, autologous stem-cell transplantation; WBRT, whole-brain radiotherapy; CR, complete response.

LOC 2023 provided molecular consensus-signature labels and BLOCAGE-01 contributed the largest trial-associated series across more than 10 French LOC centers; together they formed the development set used for coefficient estimation. BCN was reserved as the external international validation cohort and contributed an independent institution, country, scanner manufacturer, and acquisition format. This design tested locked feature definitions across NDPI and MRXS images, while the explicit patient/slide distinction prevented additional sections from being misreported as additional participants.

### 2.2 Exploratory WSI survival learning

An attention-based Cox MIL model was trained separately within each cohort. Apparent concordance was corrected with patient-level bootstrap resampling (Figure 3). Bootstrap-corrected Harrell concordance indices were 0.797 for LOC 2023, 0.797 for BLOCAGE-01, and 0.834 for BCN, consistently demonstrating outcome-related information in WSI morphology. Architecture, loss, concordance trajectories, and bootstrap specifications are reported in the Supplementary methods.

**Fig. 3.**
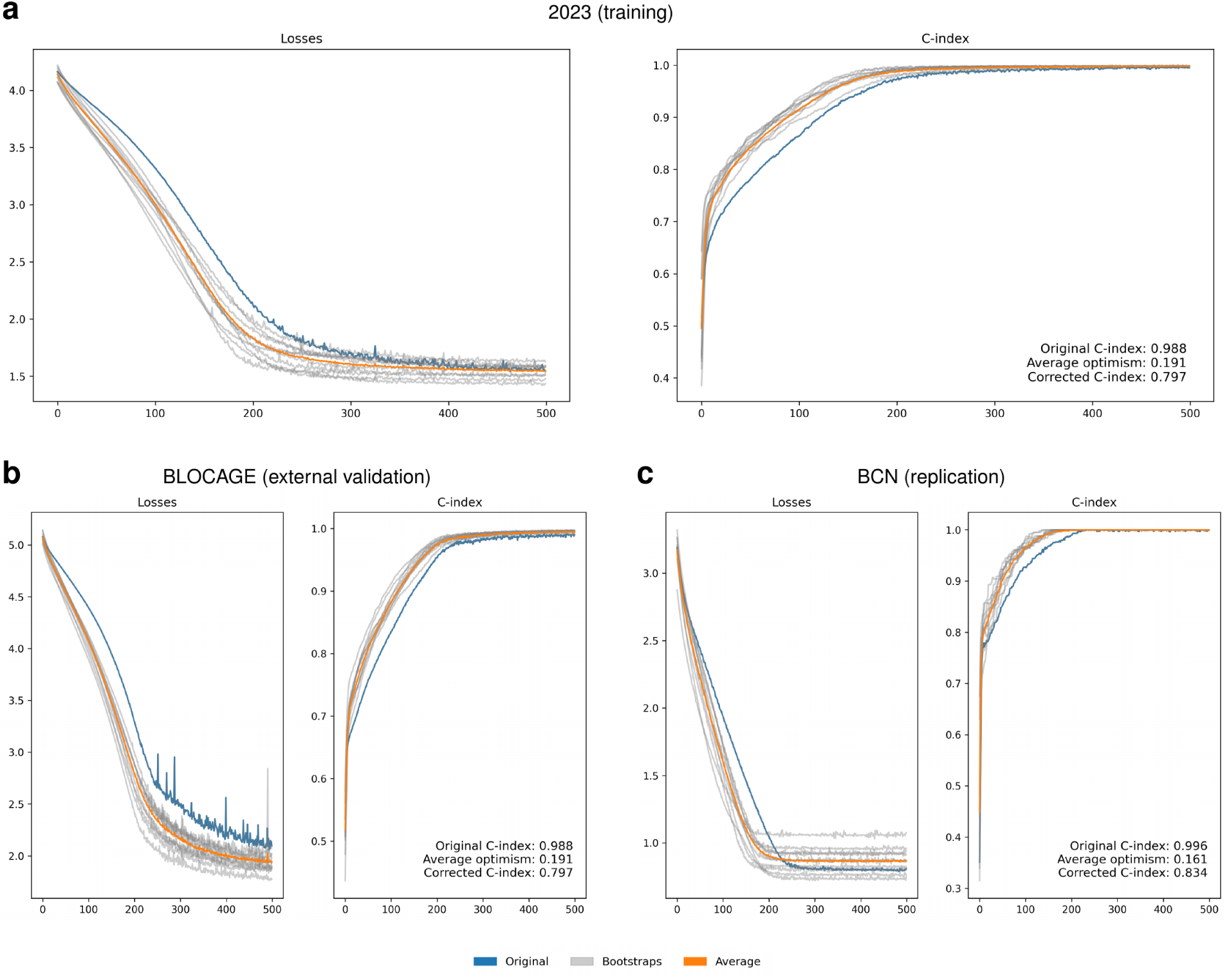
Within-cohort attention-based Cox multiple-instance-learning survival models with patient-level bootstrap correction. **a–c)** Training loss (left) and Harrell concordance-index (right) trajectories for LOC 2023 (**a**), BLOCAGE-01 (**b**), and BCN (**c**). Blue, model fitted to the original cohort; gray, each of ten patient-level bootstrap resamples evaluated in-bag; orange, mean bootstrap trajectory. Annotations report apparent concordance, mean bootstrap optimism, and optimism-corrected concordance.

Training losses decreased smoothly and apparent concordance increased in every cohort. The bootstrap correction reduced optimism while preserving concordance of approximately 0.80 (0.797– 0.834) in each dataset. These results establish that slide morphology contains reproducible outcome-related structure and motivate the subsequent feature-based analysis in which composition and spatial organization can be inspected directly.

### 2.3 An interpretable multifeature morphologic ITH score

The feature bank described slide-level prototype composition, local mixing, spatial autocorrelation, diversity, and multiscale persistence. Shannon entropy was *H* = *−*Σ*_k_ f_k_* log *f_k_* and Simpson diversity was 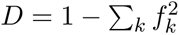 [39, 40]. Moran’s *I* measured spatial autocorrelation [41]. TP-HPM was defined as Tumor-Prototype Heterogeneity Persistence Mass, the normalized area under a multiscale Jensen– Shannon heterogeneity curve.

Eleven features spanning composition, prototype support, topology, autocorrelation, and multiscale persistence entered the elastic-net Cox model. Coefficients were estimated on pooled LOC 2023 and BLOCAGE-01 data with cohort-stratified baseline hazards. Their mathematical definitions, coefficients, scaling, and conditional interpretation are reported in the Supplementary methods.

The retained variables spanned distinct mathematical families. Prototype fractions and the dominant-prototype code summarized composition; `q_bar_0` summarized average prototype support; Moran features quantified whether assignments formed spatially coherent regions; neighborhood degree and mixing described local tissue topology; and TP-HPM integrated five-state Jensen–Shannon heterogeneity after smoothing over increasing physical scales. The simultaneous retention of composition and autocorrelation indicates that two slides with similar global prototype fractions need not receive the same score if one is spatially segregated and the other is finely intermingled.

The fitted coefficient pattern further showed that adverse risk was distributed across several morphologic axes rather than driven by a single proxy. Positive contributions included prototype-2 and prototype-3 fractions, prototype-specific and mean Moran autocorrelation, graph degree, and average prototype-0 support; spatial-mixing status, the dominant-prototype code, and five-state persistence contributed with negative conditional coefficients. Because the score is additive on the log-hazard scale, each slide can be decomposed into feature-level contributions. The exact equations, transformations, coefficients, and mathematical interpretations are provided in the Supplementary methods.

### 2.4 Association with overall survival

Continuous ITH-C was associated with OS in all three cohorts (Table 2; Figure 4). Square brackets indicate the interquartile range (IQR). In LOC 2023, lower- and higher-score groups had median OS of 61.8 months [13.9–121.2] and 15.2 months [5.7–104.7], respectively. In BLOCAGE-01, median OS was not reached [18.7–not reached] in the lower group and was 59.6 months [11.4–107.1] in the higher group. In BCN, corresponding medians were 30.2 months [5.0–82.4] and 3.6 months [1.9–29.0].

**Fig. 4.**
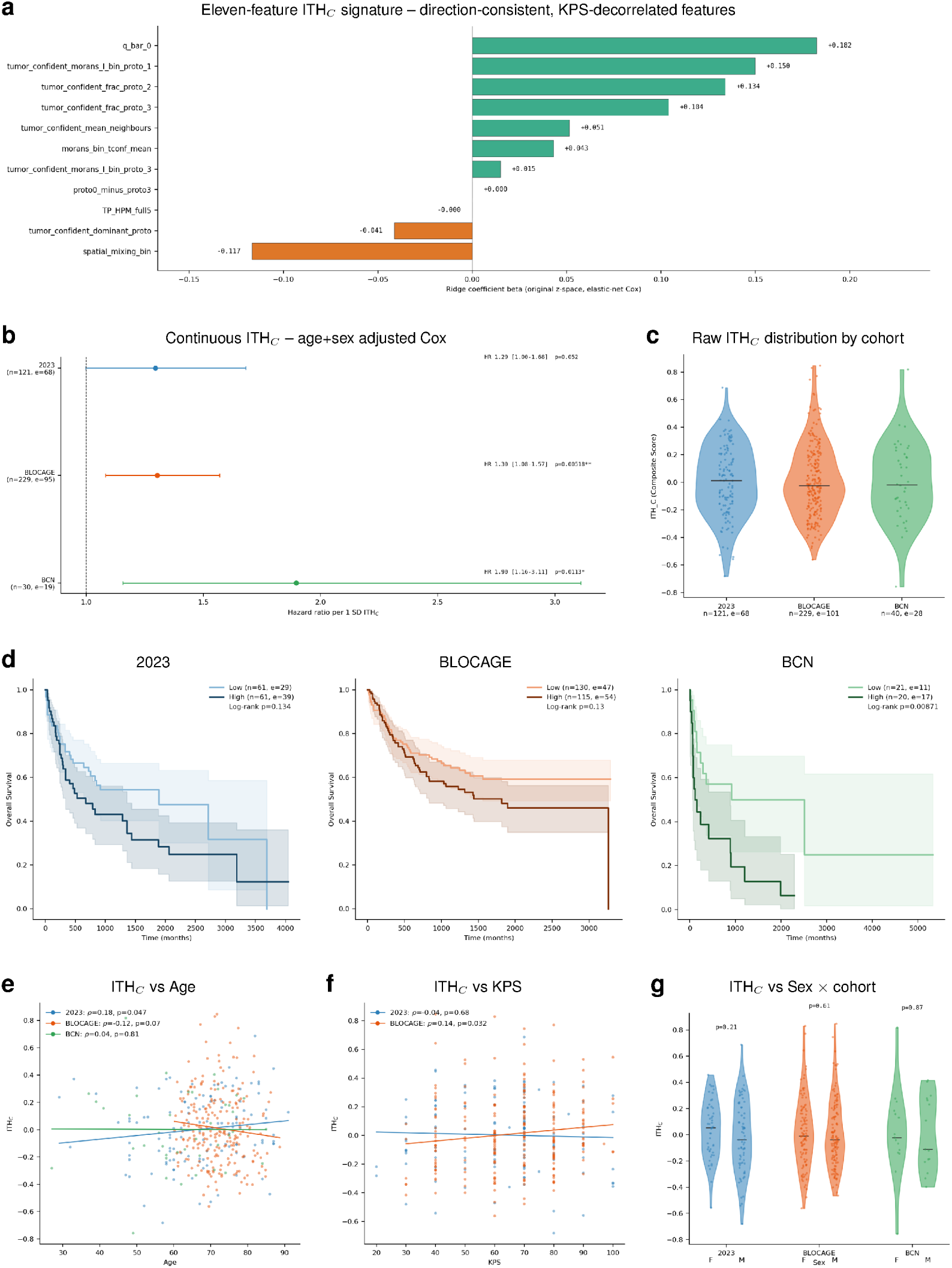
The multifeature morphologic intratumoral heterogeneity score (ITH-C) and its relationship to clinical variables and survival. **a)** Standardized elastic-net Cox coefficients for the eleven retained heterogeneity features. **b)** Cohort-specific age-, sex-, and KPS-adjusted HRs for continuous ITH-C, shown without statistical pooling. LOC 2023 and BLOCAGE-01 are development cohorts; BCN is external validation. **c)** Raw ITH-C distribution per cohort. **d)** Kaplan–Meier overall survival by median-split ITH-C with log-rank *P* values. **e,f)** ITH-C versus age (**e**) and KPS (**f**), with Spearman correlation and linear fit. **g)** ITH-C by sex within each cohort.

**Table 2:**
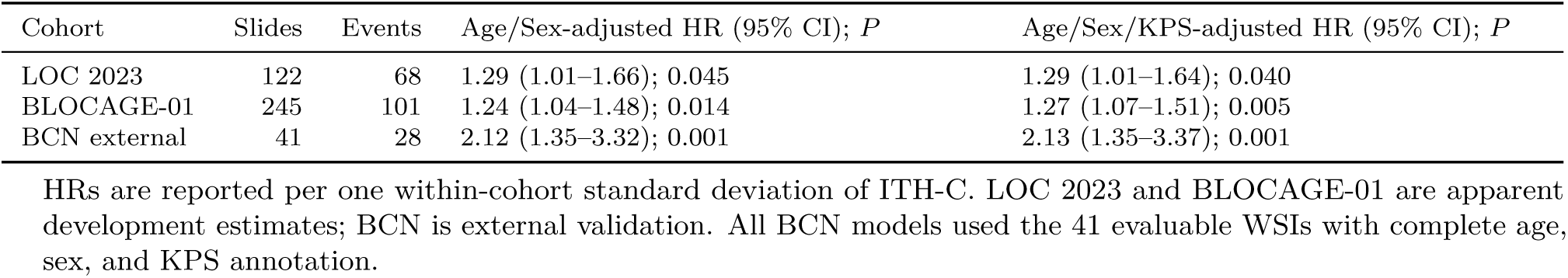
Continuous morphologic ITH score and overall survival.

| Cohort | Slides | Events | Age/Sex-adjusted HR (95% CI); $P$ | Age/Sex/KPS-adjusted HR (95% CI); $P$ |
| --- | --- | --- | --- | --- |
| LOC 2023 | 122 | 68 | 1.29 (1.01–1.66); 0.045 | 1.29 (1.01–1.64); 0.040 |
| BLOCAGE-01 | 245 | 101 | 1.24 (1.04–1.48); 0.014 | 1.27 (1.07–1.51); 0.005 |
| BCN external | 41 | 28 | 2.12 (1.35–3.32); 0.001 | 2.13 (1.35–3.37); 0.001 |

HRs are reported per one within-cohort standard deviation of ITH-C. LOC 2023 and BLOCAGE-01 are apparent development estimates; BCN is external validation. All BCN models used the 41 evaluable WSIs with complete age, sex, and KPS annotation.

LOC 2023 and BLOCAGE-01 contributed to score development, so their HRs and discrimination estimates are apparent; BCN provides the external estimate. After adjustment for age, sex, and continuous KPS, the per-SD HR was 1.29 (95% CI, 1.01–1.64) in LOC 2023, 1.27 (1.07–1.51) in BLOCAGE-01, and 2.13 (1.35–3.37) in BCN. Adding ITH-C to MSKCC increased Harrell’s *C* from 0.671 to 0.717 (Δ*C* = 0.046, bootstrap 95% CI, 0.006–0.095; likelihood-ratio *P* = 0.027), from 0.560 to 0.593 (Δ*C* = 0.033, 0.002–0.085; *P* = 0.008), and from 0.588 to 0.706 (Δ*C* = 0.118, 0.017–0.205; *P* = 0.003), respectively. For age, sex, and continuous KPS, adding ITH-C increased *C* from 0.748 to 0.754, 0.597 to 0.619, and 0.696 to 0.741; all nested likelihood-ratio tests were significant. Repeated patient-grouped downstream cross-validation retained mean MSKCC Δ*C* values of 0.040, 0.026, and 0.122. Adjusted top-versus bottom-tertile HRs were 1.97 (1.01–3.84), 1.78 (1.09–2.92), and 4.75 (1.60–14.11). Full model comparisons are reported in the Supplementary methods.

### 2.5 Spatial transcriptomic corroboration

Eight treatment-naive PCNSL Visium sections were analyzed: four from GSE203552 [26] and four from GSE230207 [42]. Starfysh [43] mapped CS1–CS4 in tissue coordinates, and the locked eleven-feature equation was projected spotwise as an RNA-based tissue readout of the H&E-derived score (Figure 5). Every biopsy contained all four programs, with regional transitions between dominant states. Because local ITH-C is a spatial projection of the same equation, its Moran autocorrelation (0.559–0.939; adjusted *P <* 0.05 in all sections) quantifies the physical organization of that projection rather than independent score validation.

**Fig. 5.**
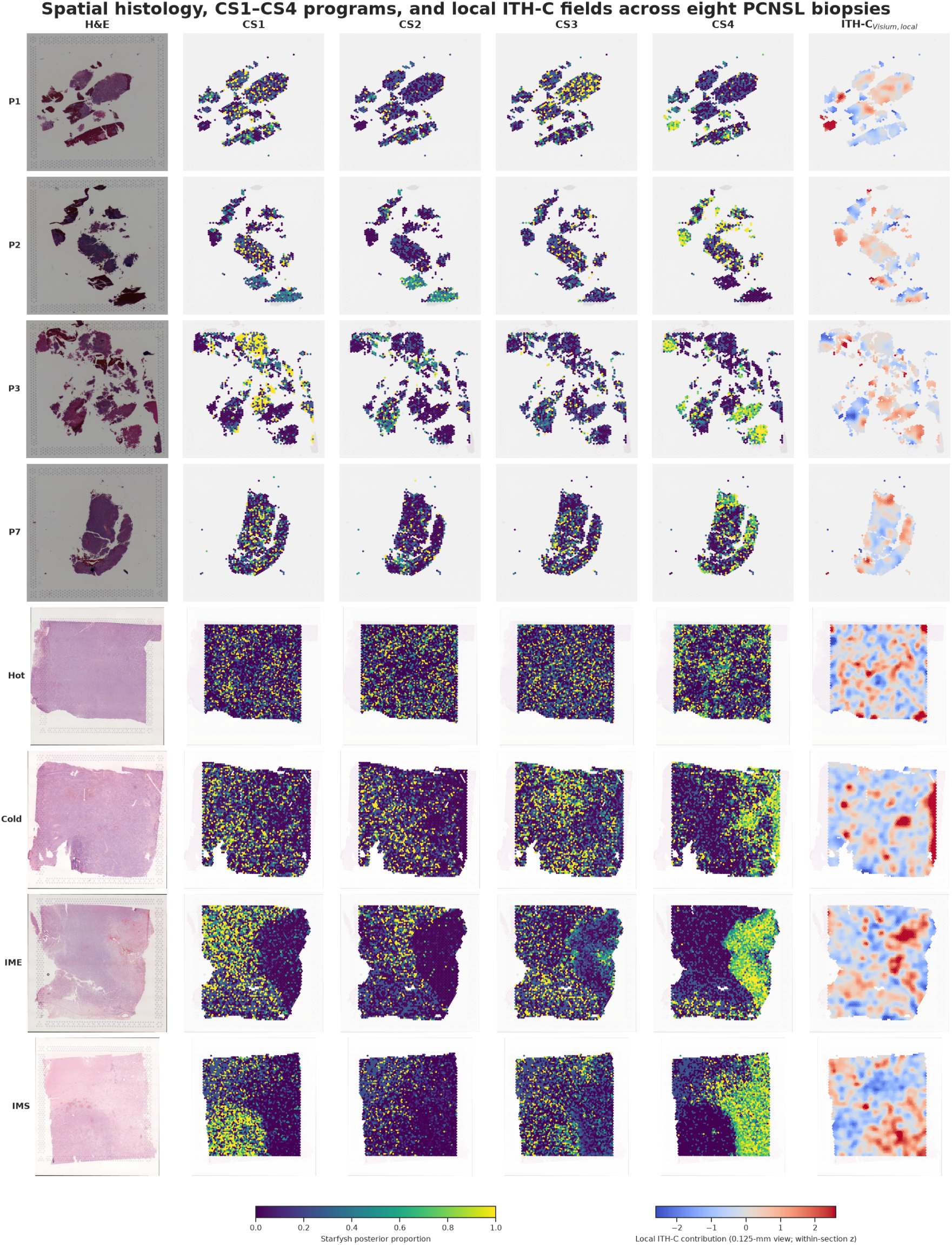
Spatial histology, CS1–CS4 programs, and local ITH-C across eight PCNSL biopsies. Rows show the H&E image, Starfysh posterior support for CS1–CS4, and the domain-adapted local ITH-C field for four GSE203552 sections (P1, P2, P3, and P7) and four GSE230207 sections (Hot, Cold, IME, and IMS). CS maps use a common 0–1 posterior scale. Local ITH-C is shown as a within-section *z* score after 0.125-mm component-aware smoothing.

The independent tissue-level finding was the association with immune programs not used to construct ITH-C. High-score regions were enriched for cytotoxic T-cell, interferon, and myeloid programs; meta-correlations were positive for cytotoxic T-cell and natural-killer enrichment and negative for the tumor/B-cell program (Supplementary Figure S6). Gain, loss, and aneuploidy formed complementary domains, while section-level aneuploidy was unrelated to mean ITH-C (*ρ* = *−*0.12, *P* = 0.779).

Median 0.125-mm Ripley enrichment was 2.281 (ITH-C), 2.546 (gain), 2.560 (loss), and 2.545 (aneuploidy), significant in all sections and decaying toward 2 mm (Figure 6). CS4 showed strongest clustering and effector, exhaustion, activation, and terminal-effector scores. Across ITH-C tertiles, these scores increased while proliferation decreased (Supplementary Tables S11–S13).

**Fig. 6.**
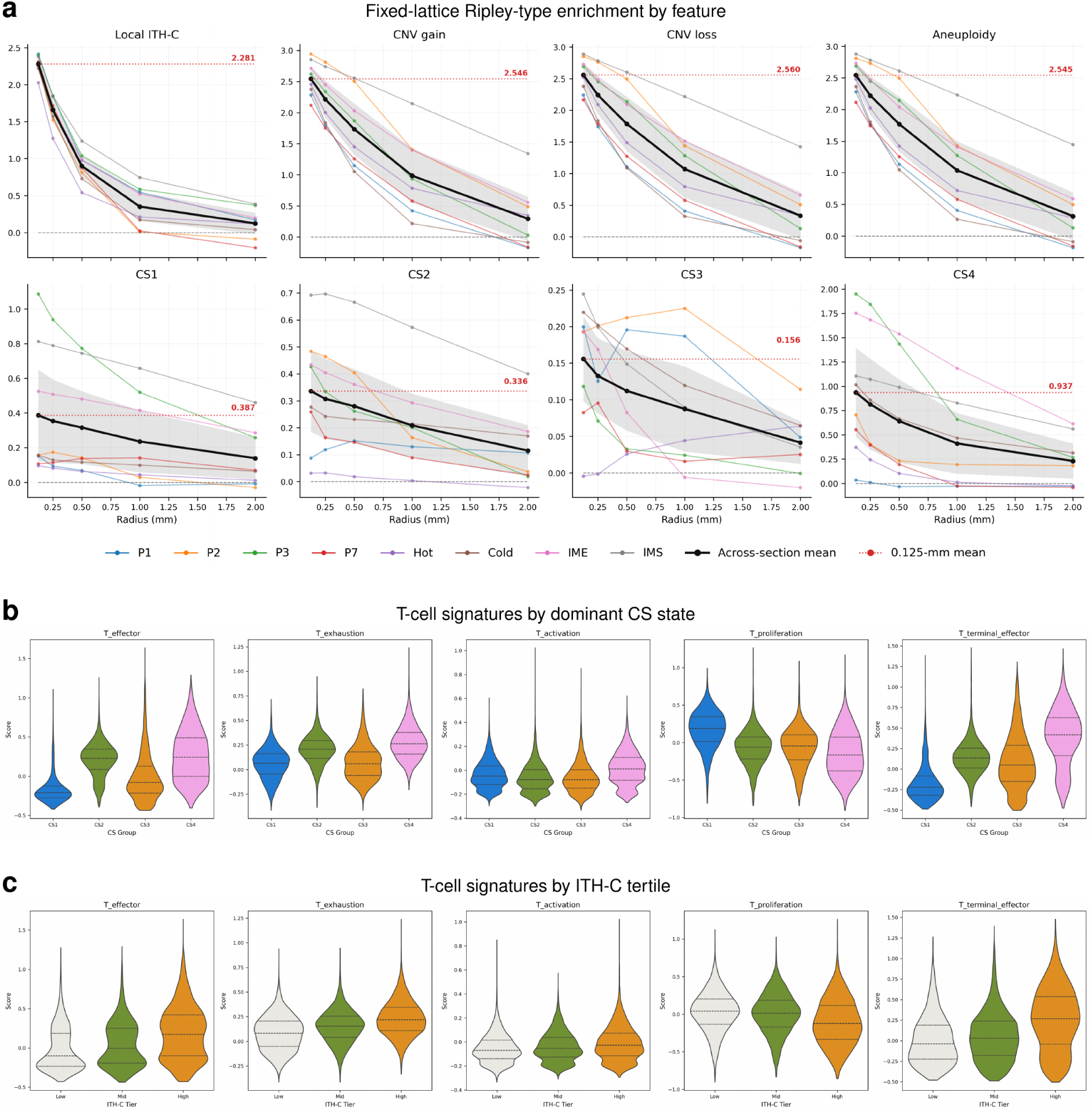
Multiscale spatial clustering and T-cell programs associated with CS1–CS4 and local ITH-C. **a)** Fixed-lattice Ripley-type enrichment of top-quartile values across 0.125–2 mm. Thin colored curves denote individual sections; black curves and gray bands denote across-section means and 95% standard-error intervals. **b)** T-cell signature distributions by dominant CS state. **c)** The same signatures across within-section low, middle, and high ITH-C tertiles. Violin-plot lines denote quartiles. RNA-inferred gain, loss, and aneuploidy are relative expression-derived fields. Signature genes and numerical summaries are provided in Supplementary Tables S11–S13.

## 3 Discussion

Across three cohorts and reference centers, routine H&E WSIs contained a consistent morphologic ITH-associated survival signal. ITH-C was associated with OS in both development cohorts and external BCN, remained significant after continuous-KPS adjustment in each cohort, and improved discrimination beyond MSKCC in all three. Bootstrap-corrected MIL concordance near 0.80 independently showed outcome-related information in slide morphology. These results position histology as a complementary prognostic source in biopsy-limited PCNSL.

The score integrates composition and organization rather than relying on a single diversity index. Prototype fractions describe which states are represented; local mixing describes adjacency; Moran’s *I* describes spatial clustering; connected components capture regional architecture; and multiscale persistence determines whether heterogeneity remains after progressive spatial smoothing. A slide with two large, separated prototype domains and a slide with the same prototype fractions intermingled at patch scale can have identical Shannon or Simpson diversity, yet differ substantially in boundary density and characteristic spatial scale. ITH-C captures these complementary dimensions and thereby operationalizes spatial ITH as a multiaxial tissue property.

This multiaxial formulation is well matched to the histopathology of PCNSL. Diagnostic biopsies can contain compact lymphoma-rich regions, angiocentric cuffs, reactive astroglial tissue, macrophage-rich interfaces, necrosis, and variable lymphoid or myeloid infiltrates within a small area. Global abundance alone cannot distinguish a coherent lymphoma compartment from dispersed perivascular or interfacial patterns. By combining abundance with adjacency and physical scale, ITH-C captures how these tissue states are assembled. The result is an integrated ecosystem phenotype that can complement clinical risk scores without requiring additional tissue consumption or a separate molecular assay.

These findings extend an emerging computational-pathology literature on tumor heterogeneity. Deep-ITH linked histology-predicted transcriptional heterogeneity to prognosis in breast cancer [20]; GBM360 connected spatial cellular architecture to glioblastoma outcome [21]; and pan-cancer histology models have predicted survival directly from WSIs [36, 37]. In large B-cell lymphoma, DLBCL-Morph demonstrated prognostic value from quantitative nuclear geometry, while spatial profiling resolved recurrent cellular neighborhoods and inflammatory niches [22–24]. The present study links H&E-derived spatial ITH with PCNSL survival, develops the score in two French series, and validates it externally in Spain.

Higher ITH-C was associated with worse OS, consistent with evolutionary models of treatment tolerance and escape [8–10]. CS1–CS4 include malignant B-cell and microenvironmental programs [25], and H&E preserves their integrated tissue manifestation. The local Visium field reused the locked feature equation, making its spatial coherence a property of the projected score; the orthogonal biological result was its association with immune expression. High-score regions showed cytotoxic, interferon, and myeloid enrichment, while CS4 combined effector, exhaustion, activation, and terminal-effector programs. Thus spatial transcriptomics connected the H&E-defined heterogeneity framework to an immune-engaged tissue state.

RNA-inferred gain, loss, and aneuploidy were spatially clustered but did not track mean ITH-C across sections, indicating complementary tissue properties. Ripley enrichment for ITH-C, CNV, and aneuploidy decayed with distance, defining their physical scales and distinguishing ITH-C from chromosomal imbalance.

The three cohorts also provide an informative natural test of clinical robustness. The molecular and clinical characterization of LOC 2023 was published by Hernández-Verdin et al. [25]. In the 107-patient image cohort analyzed here, treatment annotation recorded CAR T-cell therapy in 21 patients (20%), low-dose whole-brain radiotherapy in 11 (10%), and autologous stem-cell transplantation in 21 (20%) during the disease course. BLOCAGE-01 included patients who fulfilled the trial randomization criteria after achieving complete response to high-dose methotrexate-based induction and were subsequently assigned to rituximab–methotrexate–temozolomide maintenance or observation [44]. This complete-response requirement enriched the cohort for induction-sensitive disease and helps explain its longer median OS. BCN was a real-life cohort collected from 2008 to 2020: 8 of 40 patients received autologous stem-cell transplantation, 8 were managed palliatively from the outset, and 4 received low-dose whole-brain radiotherapy. This clinical heterogeneity may partly explain its broader and poorer survival distribution and makes the independent external association particularly informative.

Despite these differences, ITH-C retained an adverse prognostic association in every dataset. Its added value was also quantitative: MSKCC plus ITH-C increased C-index by 0.046 in LOC 2023, 0.033 in BLOCAGE-01, and 0.118 in external BCN, with significant likelihood-ratio tests; downstream patient-grouped cross-validation retained gains of 0.040, 0.026, and 0.122. The association transferred from Hamamatsu NDPI images acquired across French centers to 3DHISTECH MRXS images acquired in Spain. External BCN validation therefore extends the signal across treatment eras, cohort-entry mechanisms, countries, centers, and acquisition systems.

The concordance improvements were greatest over MSKCC, while smaller but consistently positive gains were observed after continuous KPS adjustment. This pattern indicates that ITH-C captures tissue information partly shared with age and performance status, yet still contributes morphologic and spatial information unavailable to either score alone.

Histopathology foundation models make this strategy scalable. UNI, Virchow, and Prov-GigaPath support learning without training a slide encoder de novo [31–33]. H&E models already infer driver mutations, microsatellite instability, and survival [34, 35, 37]; BRIDGE also connects histology with transcriptomic response phenotypes [45]. ITH-C complements end-to-end models by converting embeddings into inspectable spatial descriptors.

Explainability is a practical strength. Penalized Cox regression identifies signed contributions from composition, topology, autocorrelation, and persistence, allowing slide-level risk differences to be traced to prototype maps. Standardized contributions therefore provide an auditable alternative to a black-box survival output.

The approach creates several clinically relevant opportunities. The observed discrimination gain establishes complementarity to MSKCC, while evaluation beyond IELSG awaits cohorts with its complete laboratory and imaging components. A fixed ITH-C implementation could identify heterogeneous biopsies for deeper molecular profiling and support trial stratification without consuming additional tissue. Its feature-level decomposition can indicate whether risk is associated predominantly with prototype composition, local intermixing, or large-scale segregation, generating testable hypotheses for paired immunohistochemical and spatial-omic studies.

The framework can extend across tumors: disease-relevant prototype maps can be summarized with the same entropy, mixing, autocorrelation, connected-region, and persistence definitions. Brain tumors are an immediate application because spatial architecture influences sampling and resistance; pan-cancer use could reveal shared heterogeneity phenotypes. Explicit millimeter-scale features enable comparison across scanners and institutions.

Development-cohort estimates are apparent because both cohorts informed feature and penalty selection. BCN externally validates the locked score with complete age, sex, and KPS annotation, while complete IELSG components were unavailable across cohorts. Downstream patient-grouped cross-validation supported discrimination gains; prospective validation can evaluate the fully locked pipeline in broader clinical populations.

In conclusion, H&E-based ITH-C was prognostic for OS after development in two cohorts and external validation in BCN. It added discrimination beyond MSKCC and remained informative across treatment, enrollment, scanner, and acquisition differences. After prospective validation and standardized deployment, ITH-C could complement clinical scores at diagnosis to refine counseling, stratify trials, and identify patients for intensified monitoring or molecular characterization without consuming additional tissue within multidisciplinary PCNSL care pathways across treatment settings.

More broadly, this study shows how foundation-model embeddings can be transformed into interpretable measures of tissue ecosystem heterogeneity. The same strategy can support spatial risk characterization in other brain tumors and at pan-cancer scale, extending digital pathology from classification of disease and molecular alterations toward reproducible, biologically grounded survival modeling.

## 4 Methods

### 4.1 Study design, populations, and clinical annotation

LOC 2023 and BLOCAGE-01 were designated development cohorts; BCN was reserved for external international validation. LOC 2023 was a multicenter French cohort containing 107 patients with 122 evaluable diagnostic WSIs; its molecular and clinical characteristics were reported by Hernández-Verdin et al. [25]. BLOCAGE-01 contained 244 patients and 245 evaluable WSIs from more than 10 French centers. It was an open-label, multicenter randomized phase III trial in older patients eligible only after complete response to high-dose methotrexate-based induction and assigned to maintenance rituximab–methotrexate–temozolomide or observation (NCT02313389) [44]. BCN contained 40 patients and 41 evaluable WSIs collected from 2008 to 2020 at Hospital Universitari de Bellvitge and Institut Català d’Oncologia. All patients were immunocompetent and HIV negative, and all tumors were EBV negative by immunohistochemistry. OS was measured from the cohort-specific clinical origin to death; living patients were censored at last follow-up. Age, sex, and KPS were harmonized in all three analytic cohorts.

The analytic unit was the WSI because patch embeddings and prototype maps were generated for each scanned section. Patient and slide counts were retained separately, and all slides from one patient remained in the same resampling partition. Continuous-score Cox analyses used slide-level outputs; BCN models used all 41 evaluable WSIs and 28 events. Reporting followed REMARK recommendations; the completed checklist accompanies the Supplementary information [46].

ONCONEUROTEK 2 (ONTv2; NCT06314607) provides the clinical-translational framework for patients followed at Pitié-Salpêtrière Hospital from 2022 onward [47].

### 4.2 Slide acquisition and preprocessing

All WSIs were acquired at 40*×*. LOC 2023 and BLOCAGE-01 slides were NDPI files scanned on Hamamatsu NanoZoomer S60 systems at approximately 0.23 *µ*m/pixel. BCN slides were MRXS files scanned on a 3DHISTECH Pannoramic Flash III at approximately 0.24 *µ*m/pixel. Tissue was detected with Otsu thresholding [48] followed by morphological closing and contour filtering. Nonoverlapping 256 *×* 256-pixel tissue patches were extracted at level 0 after foreground and artifact filtering. The same target physical resolution and preprocessing policy was applied across centers. A representative mask is provided as Supplementary Figure S1.

The tissue mask determined eligible patch coordinates while retaining immune, stromal, vascular, necrotic, and reactive regions inside the tissue contour, consistent with measurement of a composite tissue phenotype. Quality-control operations and conversion from native pixels to physical distances are specified in the Supplementary methods.

Native patch centroids were converted to millimeters using scanner metadata before construction of spatial graphs and multiscale kernels. This ensured that a neighborhood or Gaussian smoothing radius represented the same tissue distance in NDPI and MRXS images. Tissue holes and disconnected fragments were retained as topologic boundaries, and smoothing was performed within connected components so that probability mass did not cross empty slide space. These physical-coordinate operations were central to reproducibility across acquisition systems.

### 4.3 Patch embeddings and multiple-instance learning

Each patch was embedded with UNI into a 1,024-dimensional feature vector [31]. A WSI was treated as an unordered bag of patch embeddings. Gated attention assigned normalized patch weights and formed a slide vector [28, 29]. For exploratory survival learning, a linear risk head was optimized with the negative Cox partial log-likelihood using pycox [49]. Harrell’s concordance index was calculated with comparable pairs [50]. Models were fit separately by cohort, with patient-level bootstrap resampling to reduce leakage between multiple slides from one patient. Full architecture, loss, training, and bootstrap details are in Supplementary Methods.

The survival MIL model is a learned attention-pooling Cox model, whereas TP-HPM is a deterministic multiscale summary computed from prototype probabilities and coordinates.

### 4.4 Prototype-learning MIL

The prototype MIL classifier learned four tumor prototypes corresponding to the four bulk-derived consensus-signature labels and one negative/background prototype. For patch embedding **h***_i_* and prototype ***π****_k_*, the rescaled distance *d_ik_* was converted to a soft assignment

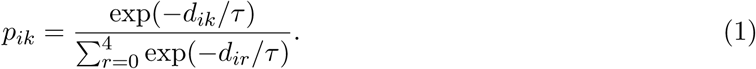

Slide-level prediction combined an attention-pooled classifier with prototype losses. Cross-validation was patient grouped. Tumor-confident patches were those with adequate positive-prototype support 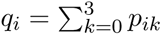. Experiments involving prototype reinitialization, repulsion, and both interventions are detailed in the Supplementary methods.

Prototype identifiers 0–3 denote the four learned CS1–CS4-associated vectors. The fifth, negative prototype represents the model’s alternative for patches with limited support from the four positive prototype states. Together, these five mutually normalized probabilities define the composition used for downstream spatial analyses.

### 4.5 Morphologic heterogeneity features

For tumor-confident patches, hard assignments were *c_i_* = arg max*_k_*_≤3_ *p_ik_* and fractions were 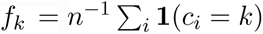. Shannon entropy and Simpson diversity were calculated as

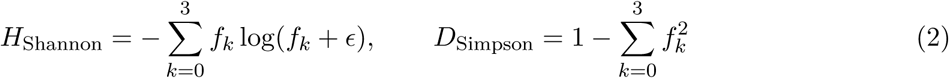

[39, 40]. A spatial graph connected neighboring retained patches. For prototype indicator *u_ik_* = **1**(*c_i_* = *k*) and weights *w_ij_*, Moran’s statistic was

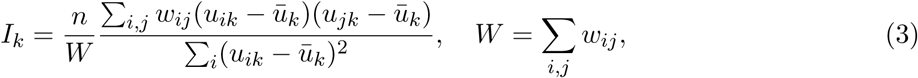

following Moran [41]. Neighborhood degree and unlike-label mixing were also recorded.

### 4.6 Tumor-Prototype Heterogeneity Persistence Mass

TP-HPM quantifies how five-state prototype heterogeneity persists under spatial smoothing. Let **p***_i_* = (*p_i_*_0_*, …, p_i_*_4_), where states 0–3 are tumor prototypes and state 4 is negative/background. For each spatial scale *ℓ ∈ {*0, 0.125, 0.25, 0.5, 1, 2*}* mm, probabilities were Gaussian-smoothed within connected tissue components. With slide mean **p̅**^(^*^ℓ^*^)^, the normalized five-state Jensen–Shannon heterogeneity was

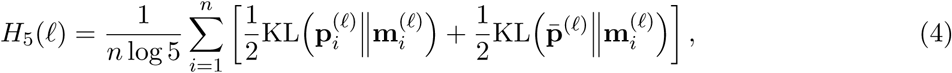

Where 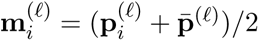. With *u*(*ℓ*) = log(1 + *ℓ/δ*) and *δ* = 0.0625 mm,

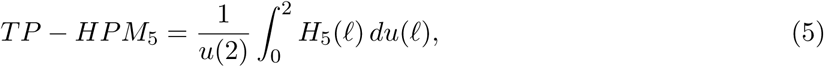

evaluated by the trapezoidal rule. The positive-prototype and positive-versus-negative decomposition, effective patch mass, characteristic radius, and fragmentation slope are derived in the Supplementary methods.

### 4.7 Penalized Cox feature combination

Candidate metrics were clustered by absolute Spearman correlation, and representative metrics were screened for KPS correlation and complementary information content. Eleven retained features were standardized within cohort:

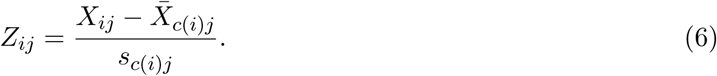

Elastic-net coefficients were estimated on pooled LOC 2023 and BLOCAGE-01 data with cohort-stratified baseline hazards. For event set *D* and risk sets *R_i_* within strata, coefficients minimized

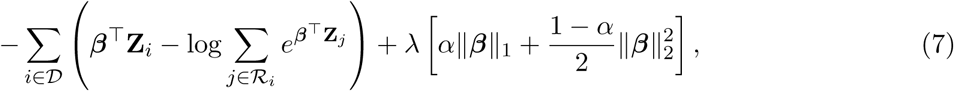

with *λ* = 0.005 and *α* = 0.5 [51–53]. The ITH-C composite was 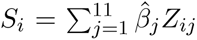. A ridge penalty of 0.001 supported downstream numerical stability. The development-derived equation was locked before external BCN evaluation. Exact definitions, transformations, and coefficients are provided in the Supplementary methods.

### 4.8 Statistical analysis

Cox models used continuous standardized ITH-C with age, sex, and continuous KPS in each cohort. Per-SD HRs used Wald 95% CIs [54], and survival summaries used Kaplan–Meier estimates [55]. Scaled Schoenfeld-residual tests supported proportional hazards in all cohorts (*P* = 0.115–0.718). Harrell’s *C* quantified discrimination [50].

MSKCC was reconstructed from age and KPS [5]. Models compared ITH-C, MSKCC, age/sex, and continuous-KPS combinations. Incremental value used Δ*C*, likelihood-ratio testing, and 24-month continuous NRI/IDI [56, 57], with 1,000 patient-cluster bootstraps and repeated fivefold patient-grouped downstream cross-validation. Cohort-specific estimates were retained without statistical pooling; adjusted tertile contrasts complemented continuous inference. Complete IELSG comparison was precluded by unavailable harmonized components. All tests were two sided; *P <* 0.05 was significant.

### 4.9 Spatial transcriptomic reanalysis

Eight treatment-naive PCNSL Visium sections were obtained from GSE203552 (P1, P2, P3, and P7) and GSE230207 (Hot, Cold, immune-microenvironment-excluded [IME], and immune-microenvironment-suppressed [IMS]) [26, 42]. Count matrices, tissue coordinates, and H&E images were barcode aligned. After expression quality control, library normalization, log transformation, and selection of 2,000 highly variable genes, CS1–CS4 marker sets were supplied as Starfysh anchors [25, 43]. Three initializations were run per section and the lowest-objective fit was retained.

Spatial projection calculated eleven ITH-C terms from Starfysh five-state fields and a six-neighbor graph within 0.13 mm. The standardized sum was component-smoothed at 0.125 mm and restandardized. Immune compartments used marker-control scores and cell2location [58]. T-cell, macrophage, and checkpoint signatures used matched controls and were compared by CS and ITH-C tertiles. infercnvpy estimated gain, loss, burden, altered-bin fraction, and aneuploidy with GENCODE v50, 100-gene windows, 10-gene steps, dynamic threshold 1.5, clipping at 3, and within-section reference. Moran’s *I* and Ripley enrichment used 199 permutations [41, 59] and Benjamini–Hochberg correction. Supplementary methods provide equations, genes, references, and section-aware analyses.

### 4.10 Software and reproducibility

Whole-slide-image processing, MIL/prototype-learning training, survival modeling, and heterogeneity-metric analysis used Python 3.10.0 with PyTorch 2.9.1, torchvision 0.24.1, OpenSlide 4.0.0 (openslide-python 1.4.3), NumPy 2.2.6, pandas 2.3.3, SciPy 1.15.3, scikit-learn 1.7.2, lifelines 0.30.0, pycox 0.3.0, matplotlib 3.10.8, and seaborn 0.13.2. Spatial analyses used the executed Python 3.12.13 environment with Starfysh 1.2.0, Scanpy 1.12.3, infercnvpy 0.6.1, PyTorch 2.13.0, pandas 2.3.3, matplotlib 3.10.9, and seaborn 0.13.2. UNI weights were obtained from https://huggingface.co/MahmoodLab/UNI. Analytic environments were version-locked to support exact computational reproducibility across platforms.

## 5 Declarations

## Data availability

Raw WSIs and individual-level clinical data cannot be publicly shared because of regulatory constraints, patient confidentiality, and the residual risk of re-identification. Qualified researchers may contact the corresponding author to discuss access under appropriate institutional, ethical, and data-transfer approvals. PCNSL Visium data are available through Gene Expression Omnibus accessions GSE203552 [26] and GSE230207 [42]. Bulk RNA-sequencing data underlying the CS1–CS4 labels are deposited at the European Genome-phenome Archive (EGA; http://www.ebi.ac.uk/ega/) under accession EGAD00001008706.

## Code availability

Complete analysis code is deposited at https://github.com/lucas-rdlr/PCNSL-ITHC and archived as a version-frozen release at Zenodo (https://doi.org/10.5281/zenodo.22180418). Trained model checkpoints for external validation, together with the corresponding derived nonidentifying analysis objects, are archived separately at Zenodo (https://doi.org/10.5281/zenodo.21886774).

## Ethics approval and consent to participate

All patients provided written informed consent to participate, in accordance with the Declaration of Helsinki. The French LOC Network data collection and processing were authorized by the Commission Nationale de l’Informatique et des Libertés (CNIL; authorization DR-2013-279) and by the relevant institutional ethics bodies. BLOCAGE-01 is registered as NCT02313389. ONCONEUROTEK 2 (ONTv2; NCT06314607) was used for patients followed at Pitié-Salpêtrière Hospital from 2022 onward. The BCN cohort study was approved by the institutional ethics committee (approval number PR356/22).

## Consent for publication

Not applicable; the manuscript contains no identifiable individual information and no third-party patient figure requiring separate publication consent.

## Acknowledgments

We thank the patients, families, clinical teams, pathology laboratories, data managers, and participating centers of the French multicenter network, BLOCAGE-01, and Hospital Universitari de Bellvitge– Institut Català d’Oncologia. We also acknowledge the HISTOMICS platform at the Paris Brain Institute (ICM), the ONCONEUROTEK brain tumor database for their support of tissue digitization, data organization, and translational research infrastructure and Biobank HUB-ICO-IDIBELL (B.0000609), integrated into the Xarxa de Bancs de Tumors de Catalunya (XBTC), for their valuable collaboration.

