## Supplementary material and figures for "Morphologic intratumoral heterogeneity from routine whole-slide histopathology is prognostic for survival in primary central nervous system lymphoma: development in the LOC Network and international external validation"

This Supplementary information reports the extended analytical definitions, sensitivity analyses, and complete clinical-model comparisons. Reporting follows the REMARK recommendations, with a completed checklist supplied as a separate Supplementary file [1].

### S1 WSI acquisition, masking, tiling, and embedding

LOC 2023 and BLOCAGE-01 WSIs were acquired at 40× with Hamamatsu NanoZoomer S60 scanners, stored as NDPI, and sampled at approximately 0.23 $\mu$m/pixel. BLOCAGE-01 represented more than 10 French centers. BCN slides were acquired at 40× with a 3DHISTECH Pannoramic Flash III, stored as MRXS, and sampled at approximately 0.24 $\mu$m/pixel.

The CLAM preprocessing framework was used [2]. Similar weakly supervised pipelines have enabled large-scale WSI learning without region-level labels [3]. In the HSV representation, Otsu's between-class variance criterion selected a tissue threshold [4]. For candidate threshold $t$, Otsu's method maximizes

$$\sigma_B^2(t) = \omega_0(t)\omega_1(t)\left[\mu_0(t) - \mu_1(t)\right]^2, \tag{S1}$$

where $\omega_0, \omega_1$ are class probabilities and $\mu_0, \mu_1$ are the corresponding mean intensities. Morphological closing removed small discontinuities, internal holes were filtered, and only retained tissue contours were tiled at level 0.

Nonoverlapping $256 \times 256$-pixel patches were retained after foreground filtering. At the native scan resolutions, a patch spans approximately 58.9 $\mu$m in the French images and 61.4 $\mu$m in BCN before any feature-extractor resizing. Patch centroids remained expressed in the WSI coordinate frame and were converted to physical distances for scale-dependent spatial operations. This conversion is essential: applying the same Gaussian width in pixels to scanners with different microns per pixel would give different biological smoothing scales.

Each RGB patch was transformed with the UNI preprocessing pipeline and encoded as a 1,024-dimensional feature vector [5]. UNI belongs to a broader generation of self-supervised pathology encoders that includes Phikon, Virchow, Prov-GigaPath, and CTransPath [6–9]. The encoder was used as a fixed representation for downstream analyses.

#### S1.1 Mask interpretation and quality control

The mask defined tissue eligible for patch extraction, not tumor confirmed by a neuropathologist. Retained tissue could therefore include lymphoma-rich regions, reactive brain, hemorrhage, necrosis, vessels, immune infiltrates, and technical artifacts not removed by the automated filters. This broad inclusion is consistent with the bulk-derived CS labels and with the goal of measuring a composite tumor-microenvironment phenotype, but it prevents interpreting every retained patch as malignant.

Slides with empty masks, unreadable pyramids, irreconcilable coordinate metadata, or no retained patches were excluded. Image-level QC recorded scanner and format, microns per pixel, tissue area, retained patch count, fraction of rejected patches, number of connected tissue components, color-distribution summaries, and thumbnails of the mask and sampled patch grid. Figure S1 illustrates the automated mask used before tiling.

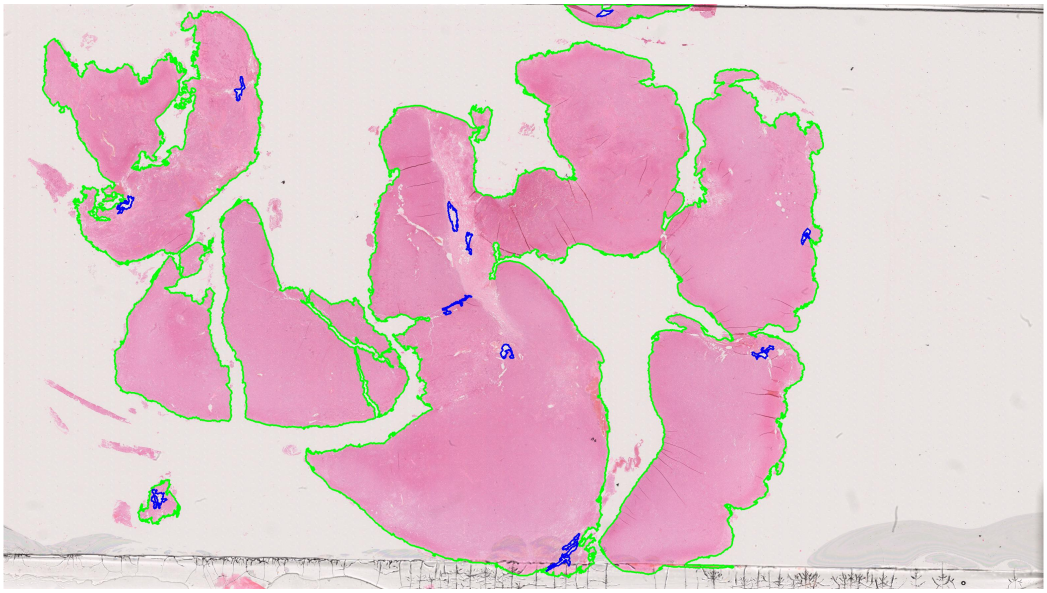

**Fig. S1. Representative automated tissue segmentation and retained tissue mask before patch extraction.** The green contour defines the tissue region submitted to tiling; internal exclusions and artifacts are removed before nonoverlapping level-0 patches are generated.

### S2 Exploratory attention-based Cox MIL

For a slide bag $\{\mathbf{h}_i\}_{i=1}^{n}$, gated attention [10] was

$$e_i = \mathbf{w}^\top \left[\tanh(\mathbf{V}\mathbf{h}_i) \odot \sigma(\mathbf{U}\mathbf{h}_i)\right], \tag{S2}$$

$$a_i = \frac{\exp(e_i)}{\sum_j \exp(e_j)}, \qquad \mathbf{z} = \sum_i a_i \mathbf{h}_i, \qquad r = \boldsymbol{\gamma}^\top \mathbf{z}. \tag{S3}$$

The negative Cox partial log-likelihood was

$$\mathcal{L}_{\text{Cox}} = -\sum_{i:\delta_i=1} \left[ r_i - \log \sum_{j:T_j \geq T_i} \exp(r_j) \right], \tag{S4}$$

implemented with pycox [11, 12]. Each cohort was modeled separately for 500 epochs. Patient-level bootstrap resampling provided the apparent concordance, estimated optimism, and subtraction-corrected concordance reported in Table S1.

#### S2.1 Bag construction and attention interpretation

For slide $s$, the bag matrix is $\mathbf{H}_s = [\mathbf{h}_{s1}, \ldots, \mathbf{h}_{sn_s}]^\top$. The attention weights satisfy $a_{si} \geq 0$ and $\sum_i a_{si} = 1$, so $\mathbf{z}_s$ is a convex combination of patch embeddings. A large $a_{si}$ means that patch $i$ contributed strongly to the fitted slide representation; it does not by itself establish that the patch is a causal adverse region. Attention can be redistributed among correlated patches and should be interpreted together with spatial maps and perturbation analyses.

The risk head produces a real-valued log-risk score. The Cox partial likelihood compares the score of each event slide with scores in its risk set and does not estimate the baseline hazard. Tied event times were handled consistently across fits. Preprocessing and batching preserved the mapping between slide, patient, event time, and embeddings despite variation in bag size.

##### S2.2 Bootstrap optimism correction

The bootstrap procedure fits the model on the original cohort and on patient-level resamples. For bootstrap sample $b$, the model is trained on resampled patients and evaluated on both the bootstrap data and the original cohort. Optimism is

$$O_b = C_b^{\text{boot}} - C_b^{\text{original}}, \qquad \bar{O} = \frac{1}{B}\sum_{b=1}^{B} O_b, \qquad C_{\text{corrected}} = C_{\text{apparent}} - \bar{O}. \tag{S5}$$

This subtraction-based correction quantifies the decrease from apparent to optimism-corrected concordance. Bootstrap correction is a standard method for internal model evaluation [13]. The resulting corrected values were 0.797, 0.797, and 0.834 across the three cohorts.

##### S2.3 Concordance and censoring

Harrell's $C$ uses comparable pairs and measures concordance between observed event ordering and predicted risk [14]. Uno's inverse-probability-of-censoring weighted concordance provides a complementary estimator at a prespecified horizon [15]. Harrell's $C$ was used consistently for the three cohort-specific models.

Table S1: Exploratory within-cohort Cox MIL results.

| Cohort | Apparent Harrell $C$ | Estimated optimism | Corrected $C$ |
|---|---|---|---|
| LOC 2023 | 0.988 | 0.191 | 0.797 |
| BLOCAGE-01 | 0.988 | 0.191 | 0.797 |
| BCN | 0.996 | 0.161 | 0.834 |

For risk scores $r_i$, Harrell's concordance index was

$$\widehat{C} = \frac{\sum_{i,j} \mathbf{1}(T_i < T_j)\delta_i \left[\mathbf{1}(r_i > r_j) + \frac{1}{2}\mathbf{1}(r_i = r_j)\right]}{\sum_{i,j} \mathbf{1}(T_i < T_j)\delta_i}, \tag{S6}$$

where only comparable pairs contribute [14].

The corrected values, consistently at or near 0.80 (0.797–0.834), demonstrate that WSI morphology carries substantial outcome-related information in each clinical setting. These cohort-specific models serve as an outcome-sensitive representation layer and support the subsequent interpretable feature analysis.

### S3 Prototype-learning MIL

#### S3.1 Architecture and loss

The prototype model used four positive tumor prototypes $\boldsymbol{\pi}_0, \ldots, \boldsymbol{\pi}_3$ and one negative/background prototype $\boldsymbol{\pi}_4$. Projected patch embeddings and prototypes were unit normalized. Let $d_{ik}$ be the model distance between patch $i$ and prototype $k$. Soft assignment at temperature $\tau$ was

$$p_{ik} = \frac{\exp(-d_{ik}/\tau)}{\sum_{r=0}^{4} \exp(-d_{ir}/\tau)}. \tag{S7}$$

The total loss combined slide-level cross-entropy, prototype guidance, and regularization:

$$\mathcal{L} = \mathcal{L}_{\mathrm{CE}} + \lambda_g \mathcal{L}_{\mathrm{guide}} + \lambda_o \|\boldsymbol{\Pi}\boldsymbol{\Pi}^\top - \mathbf{I}\|_F^2. \tag{S8}$$

The four-class target was the bulk-derived CS1–CS4 label from LOC 2023 [16]. Because those labels reflect bulk tumor and microenvironmental RNA, a patch prototype is a latent tissue state and not a purified malignant-cell state.

Prototype learning follows the general principle of representing observations by proximity to learned reference vectors [17, 18]. It is also related to vector-quantized representation learning, in which unused or crowded code vectors can create codebook collapse [19]. The model-specific distances were converted to assignments only after the checkpoint was fixed for that fold. Prototype numbers are nominal: prototype 0 is not intrinsically lower or biologically earlier than prototype 3.

#### S3.2 Positive and negative prototype roles

The four positive prototypes were aligned with the four slide-level consensus labels during training. The fifth prototype served as a negative/background alternative. Let

$$q_i = \sum_{k=0}^{3} p_{ik} = 1 - p_{i4} \tag{S9}$$

denote total positive support. Conditional positive probabilities are $r_{ik} = p_{ik}/(q_i + \epsilon)$. A patch can therefore have a relatively clear positive composition $\mathbf{r}_i$ but low total positive support $q_i$. Retaining both quantities distinguishes uncertainty about tumor-versus-background status from uncertainty among the four positive states.

The negative prototype was not trained from a pathologist-curated normal-brain atlas. It should therefore be read as the model's residual or alternative tissue prototype, not as a definitive label for non-neoplastic brain. Similarly, positive assignments should not be interpreted as local transcriptomic CS calls without paired spatial validation.

#### S3.3 Classification and guidance objectives

The slide-level cross-entropy term optimizes prediction of the bulk consensus label. The guidance term encourages patch-prototype relations to follow the model's attention-derived pseudo-targets, while the orthogonality term discourages identical prototype vectors. These objectives need not optimize the same geometry. A classifier can improve by attending to a small discriminative region even if global nearest-prototype assignments remain unbalanced. This explains why above-chance slide classification can coexist with rare use of two prototypes.

##### S3.4 Cross-validation and classification results

Patient-grouped 10-fold cross-validation produced out-of-fold classification estimates. Baseline mean one-versus-rest macro-AUC was 0.745 (range, 0.521–0.878) and accuracy was 0.507 (range, 0.259–0.733) versus a nominal four-class chance level of 0.25 (Table S2). Two positive prototypes received few nearest-prototype patches in several folds, motivating prototype-repulsion and dead-prototype-reinitialization experiments.

All slides from the same patient were assigned to the same fold. Performance was calculated only from held-out fold predictions and summarized across folds. Macro-AUC averaged one-versus-rest AUCs so that every class contributed equally. Accuracy retained the observed class distribution and was compared with the nominal balanced-class chance level only as orientation; it was not used as a formal null model.

Table S2: Out-of-fold prototype-classification performance.

| Configuration | Mean AUC (range) | Mean accuracy (range) | Improved/worse/tied folds | $P$ vs baseline |
|---|---|---|---|---|
| Baseline | 0.745 (0.521–0.878) | 0.507 (0.259–0.733) | – | – |
| Repulsion | 0.745 (0.521–0.878) | 0.507 (0.259–0.733) | 0/0/10 | 1.000 |
| Dead-prototype reinitialization | 0.736 (0.521–0.903) | 0.520 (0.259–0.667) | 3/4/3 | 0.578 |
| Combined | 0.739 (0.521–0.908) | 0.527 (0.259–0.633) | 4/3/3 | 0.375 |

For repulsion margin $m$, the additional loss was

$$\mathcal{L}_{\text{rep}} = \sum_{k<l} \max\{0, m - \|\boldsymbol{\pi}_k - \boldsymbol{\pi}_l\|_2\}^2. \tag{S10}$$

For dead-prototype reinitialization, a prototype with nearest-assignment share below the prespecified threshold was replaced at the scheduled update by a pooled observed patch embedding. The combined configuration increased use of the most underrepresented prototype, while overall classification accuracy remained similar (paired Wilcoxon $P = 0.375$).

Repulsion changes the geometry continuously by penalizing prototype pairs closer than margin $m$. Reinitialization is discontinuous: a low-use prototype is replaced with an observed embedding and then resumes gradient updates. The combined strategy addresses both crowding and dead prototypes. However, increasing prototype use is not sufficient evidence of improved representation; a reseeded prototype may simply partition common morphology without gaining molecular or prognostic meaning.

Fold-to-fold variation was quantified with maps, prototype shares, and region features. The downstream analysis therefore integrated feature families across composition, confidence, topology, autocorrelation, and multiscale persistence rather than relying on one visually selected patch map.

##### S3.5 Prototype learning was informative but unstable

The prototype-learning MIL model predicted the four bulk-derived consensus signatures above chance, although two tumor prototypes were infrequently selected by nearest-prototype assignment and fold-to-fold maps varied. Training modifications improved prototype use but did not consistently improve classification. Detailed accuracies, AUCs, mitigation experiments, and single-feature survival tests are reported in the Supplementary methods. Figure S2 illustrates the fold-dependent prototype maps.

The discrepancy between slide-level classification and patch-level prototype use is important. A discriminative attention head can classify a WSI from a limited subset of patches even when the global prototype geometry is poorly balanced. Conversely, a prototype may be used rarely because it represents a narrow but informative morphologic state, because another prototype occupies a larger basin in embedding space, or because optimization has produced partial prototype collapse. The fold-specific maps and the lack of a consistent accuracy improvement after repulsion or reinitialization favored the latter two technical explanations over a simple biological interpretation.

CS1–CS4 were learned from bulk RNA profiles and therefore incorporate malignant-cell and microenvironmental information [16]. H&E captures the same integrated tissue ecosystem, including lymphoma-cell morphology, immune and stromal organization, vascular structures, and reactive compartments. The prototype maps consequently represent composite morphologic and microenvironmental states, a biologically relevant level for spatial ITH analysis.

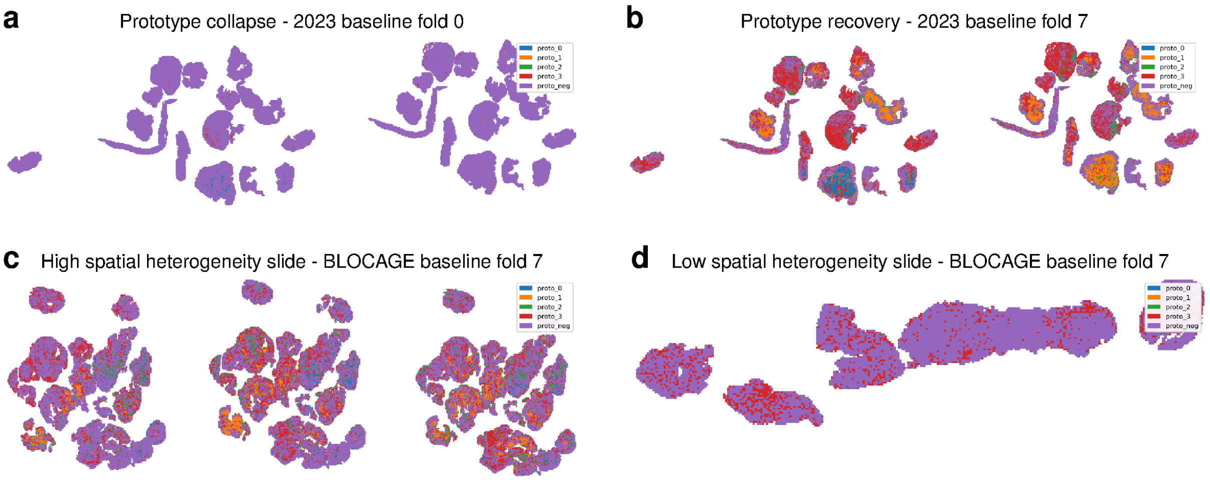

**Fig. S2. Fold-dependent prototype maps and contrasting spatial heterogeneity. a,b)** The same representative LOC 2023 WSI scored by two baseline TPMIL fold checkpoints. In fold 0 (**a**), the under-represented tumor prototype was the nearest assignment for 33 of 30,376 patches (0.1%); in fold 7 (**b**), it occupied 3,807 patches (12.5%) and formed a contiguous region of 918 patches. **c,d)** Representative BLOCAGE-01 WSIs with high (**c**; largest contiguous region, 217 patches; 4.8% of tissue) and low (**d**; largest contiguous region, 0 patches) extent of this prototype, both scored with the fold-7 checkpoint.

#### S4 Patch assignment strategies and heterogeneity feature bank

Three assignment strategies were evaluated: (1) full five-state competition; (2) tumor-only assignment to the nearest of prototypes 0–3; and (3) tumor-confident assignment, retaining only patches whose positive support exceeded the negative/background support criterion. Let

$$q_i = \sum_{k=0}^{3} p_{ik}, \qquad r_{ik} = \frac{p_{ik}}{q_i + \epsilon}, \quad k = 0, \ldots, 3. \tag{S11}$$

Here $q_i$ is tumor-prototype support and $\mathbf{r}_i$ is the conditional composition among positive prototypes. Hard positive assignment was $c_i = \arg\max_{k \le 3} r_{ik}$.

The full strategy preserves the original competition between all five prototypes. The tumor-only strategy forces every retained patch to one of the four positive prototypes and is therefore sensitive to background tissue that resembles a positive state more than the other positive alternatives. The tumor-confident strategy avoids this reassignment by excluding patches that do not satisfy the positive-support criterion. Agreement between full and tumor-confident features can indicate robustness to background handling, but does not constitute an independent validation because both derive from the same prototype distances.

##### S4.1 Composition and diversity

With $n_k = \sum_i \mathbf{1}(c_i = k)$ and $f_k = n_k / \sum_l n_l$, the dominant prototype was $\arg\max_k f_k$, and `proto0_minus_proto3` was $f_0 - f_3$. Shannon entropy and Simpson diversity were

$$H = -\sum_{k=0}^{3} f_k \log(f_k + \epsilon), \qquad H_{\text{norm}} = \frac{H}{\log 4}, \qquad D = 1 - \sum_{k=0}^{3} f_k^2 \tag{S12}$$

[20, 21]. The effective prototype number was $\exp(H)$.

Entropy is maximized when the four fractions are equal and is zero when one prototype occupies all assigned patches. Simpson diversity is the probability that two independently sampled assigned patches have different labels. Neither statistic uses coordinates. Their role is to quantify global composition before spatial structure is introduced.

##### S4.2 Neighborhood mixing and connected regions

The spatial graph connected retained grid neighbors within the configured adjacency radius. Degree was $d_i = \sum_j w_{ij}$. Unlike-label mixing was

$$M = \frac{\sum_{i,j} w_{ij} \mathbf{1}(c_i \neq c_j)}{\sum_{i,j} w_{ij}}, \tag{S13}$$

and same-label neighborhood support was

$$N_{\text{same}} = \frac{1}{n} \sum_i \frac{\sum_j w_{ij} \mathbf{1}(c_i = c_j)}{d_i}. \tag{S14}$$

Connected-component features included the number of components per prototype, maximum component size, mean component size, component-size dispersion, and boundary density.

For prototype $k$, connected components are maximal node sets joined by paths of label $k$. If their sizes are $A_{k1}, \dots, A_{kR_k}$, the maximum region is $\max_r A_{kr}$ and the mean is $R_k^{-1} \sum_r A_{kr}$. Boundary density is the proportion of graph edges that connect different hard labels:

$$B = \frac{\sum_{i<j} w_{ij} \mathbf{1}(c_i \neq c_j)}{\sum_{i<j} w_{ij}}. \tag{S15}$$

High $B$ or $M$ indicates fine interdigitation, whereas a large maximum component and high positive Moran's $I$ indicate spatial segregation. These measures are complementary rather than redundant in all slides.

##### S4.3 Spatial autocorrelation

For $u_{ik} = \mathbf{1}(c_i = k)$, $\bar{u}_k = n^{-1} \sum_i u_{ik}$, and $W = \sum_{i,j} w_{ij}$,

$$I_k = \frac{n}{W} \frac{\sum_{i,j} w_{ij}(u_{ik} - \bar{u}_k)(u_{jk} - \bar{u}_k)}{\sum_i (u_{ik} - \bar{u}_k)^2 + \epsilon} \tag{S16}$$

was Moran's $I$ [22]. Its mean across the four positive prototypes was also retained. Geary's statistic was

$$C_k = \frac{n-1}{2W} \frac{\sum_{i,j} w_{ij}(u_{ik} - u_{jk})^2}{\sum_i (u_{ik} - \bar{u}_k)^2 + \epsilon} \tag{S17}$$

[23].

Moran's $I$ compares cross-products of deviations and is sensitive to broad spatial autocorrelation. Geary's $C$ uses squared local differences and is more sensitive to short-range boundaries. Under common conventions, positive autocorrelation corresponds to $I > 0$ and $C < 1$, but finite irregular graphs and rare labels can shift their empirical ranges. When a prototype is absent or constant, the variance denominator vanishes; that prototype-specific statistic is undefined and should not be replaced with a biologically meaningful zero.

##### S4.4 Physical scale and graph construction

Spatial neighbors were defined from patch centroids after accounting for the scan's microns-per-pixel metadata. The graph is computed within retained tissue, so missing patches and holes can reduce degree at tissue boundaries. Mean degree therefore partly reflects tissue geometry and mask fragmentation. It was retained in the feature bank because this geometry can interact with region metrics, but its interpretation is different from same-label neighbor support.

The feature bank deliberately included global, local, and multiscale quantities. Global composition cannot distinguish rearrangements; a single graph radius can miss structures larger than one neighborhood; and multiscale smoothing alone can obscure which prototype contributes. Combining these families allows the penalized model to select complementary axes, while correlation reduction limits multiple nearly identical proxies.

##### S4.5 Correlation screening and degeneracy handling

Pairwise Spearman correlation was used because many region metrics are skewed and contain ties. Metrics with excessive degeneracy, no variation, or insufficient finite observations were excluded from the corresponding comparison. Finite fold-level outputs were averaged after confirming that the feature definition was constant across folds. The complete feature families are summarized in Table S3.

Table S3: Feature families calculated before correlation reduction.

| Family | Examples and interpretation |
|---|---|
| Prototype support | $\bar{q} = n^{-1}\sum_i q_i$; $\bar{q}_0 = n^{-1}\sum_i p_{i0}$; effective positive patch mass $\sum_i q_i$ |
| Composition | $f_k$; $f_0 - f_3$; dominant prototype; Shannon entropy; normalized entropy; Simpson diversity |
| Neighborhood | Mean degree; same-label neighbor fraction; unlike-label mixing; boundary density |
| Regions | Component count; largest component; mean component size; component-size variability |
| Autocorrelation | Prototype-specific and mean Moran's $I$; prototype-specific Geary's $C$ |
| Persistence | Five-state, positive-conditional, and positive-versus-negative TP-HPM; $r_{50}$; initial fragmentation slope |

#### S5 Tumor-Prototype Heterogeneity Persistence Mass

##### S5.1 Probability field and spatial smoothing

TP-HPM means Tumor-Prototype Heterogeneity Persistence Mass. The `tp_hpm` function receives the patch-to-prototype distance matrix, level-0 patch coordinates, physical pixel size, temperature $\tau$, and the prespecified smoothing grid. Distances are rescaled to a common numerical range and converted to five-state probabilities with softmax $(-d/\tau)$.

Let $\widetilde{d}_{ik}$ be the rescaled distance. The unsmoothed five-state probability vector is

$$\mathbf{p}_i^{(0)} = \mathrm{softmax}\left(-\widetilde{d}_{i0}/\tau, \ldots, -\widetilde{d}_{i4}/\tau\right). \tag{S18}$$

Temperature controls assignment sharpness. Small $\tau$ approaches hard nearest-prototype assignment, whereas large $\tau$ produces smoother probability distributions. Jensen–Shannon heterogeneity is calculated from the full probability vectors at the prespecified temperature, while alternative temperatures quantify sensitivity to assignment sharpness.

For scale $\ell$, each probability channel was smoothed with a Gaussian kernel within each connected tissue component:

$$p_{ik}^{(\ell)} = \frac{\sum_{j\in C(i)} K_\ell(\mathbf{r}_i - \mathbf{r}_j)p_{jk}}{\sum_{j\in C(i)} K_\ell(\mathbf{r}_i - \mathbf{r}_j) + \epsilon}, \quad K_\ell(\mathbf{r}) = \exp\left(-\frac{\|\mathbf{r}\|_2^2}{2\ell^2}\right). \tag{S19}$$

The analyzed scales were $\mathcal{L} = \{0, 0.125, 0.25, 0.5, 1, 2\}$ mm. Smoothing within $C(i)$ prevents probability mass from crossing disconnected tissue gaps.

Masked normalization in the denominator corrects edge attenuation. Without this term, a patch near a tissue boundary would receive less total kernel mass and its probability vector could shrink solely because neighboring coordinates are absent. Each smoothed vector was renormalized to the probability simplex within numerical tolerance. At $\ell = 0$, the original unsmoothed probabilities were used rather than a singular Gaussian.

#### S5.2 Weighted Jensen–Shannon decomposition

For distributions $\mathbf{a}, \mathbf{b}$ and midpoint $\mathbf{m} = (\mathbf{a} + \mathbf{b})/2$, define

$$\mathrm{JS}(\mathbf{a}, \mathbf{b}) = \tfrac{1}{2} \sum_k a_k \log \frac{a_k + \epsilon}{m_k + \epsilon} + \tfrac{1}{2} \sum_k b_k \log \frac{b_k + \epsilon}{m_k + \epsilon}. \tag{S20}$$

At scale $\ell$, the slide mean is $\bar{\mathbf{p}}^{(\ell)} = n^{-1} \sum_i \mathbf{p}_i^{(\ell)}$. Full five-state heterogeneity is

$$J_5(\ell) = \frac{1}{n} \sum_i \mathrm{JS}\Big(\mathbf{p}_i^{(\ell)}, \bar{\mathbf{p}}^{(\ell)}\Big), \qquad H_5(\ell) = \frac{J_5(\ell)}{\log 5}. \tag{S21}$$

Let $q_i^{(\ell)} = \sum_{k=0}^{3} p_{ik}^{(\ell)}$, $\bar{q}^{(\ell)} = n^{-1} \sum_i q_i^{(\ell)}$, and $\mathbf{r}_i^{(\ell)} = (p_{i0}^{(\ell)}, \ldots, p_{i3}^{(\ell)})/(q_i^{(\ell)} + \epsilon)$. Positive-conditional heterogeneity uses $q_i$ as mass:

$$J_+(\ell) = \frac{\sum_i q_i^{(\ell)} \mathrm{JS}\Big(\mathbf{r}_i^{(\ell)}, \bar{\mathbf{r}}_q^{(\ell)}\Big)}{\sum_i q_i^{(\ell)} + \epsilon}, \quad \bar{\mathbf{r}}_q^{(\ell)} = \frac{\sum_i q_i^{(\ell)} \mathbf{r}_i^{(\ell)}}{\sum_i q_i^{(\ell)} + \epsilon}, \quad H_+(\ell) = \frac{J_+(\ell)}{\log 4}. \tag{S22}$$

Positive-versus-negative heterogeneity uses the binary field $\mathbf{b}_i^{(\ell)} = (q_i^{(\ell)}, 1 - q_i^{(\ell)})$:

$$J_\pm(\ell) = \frac{1}{n} \sum_i \mathrm{JS}\Big(\mathbf{b}_i^{(\ell)}, (\bar{q}^{(\ell)}, 1 - \bar{q}^{(\ell)})\Big), \qquad H_\pm(\ell) = \frac{J_\pm(\ell)}{\log 2}. \tag{S23}$$

The implementation's decomposition is

$$J_5(\ell) = J_\pm(\ell) + \bar{q}^{(\ell)} J_+(\ell), \tag{S24}$$

up to numerical tolerance. This equation separates spatial variation in tumor-versus-background support from variation among the four tumor prototypes.

The weights $q_i^{(\ell)}$ in $J_+(\ell)$ ensure that patches with minimal positive support contribute little to heterogeneity among positive prototypes. The binary term $J_\pm$ answers a different question: whether tumor-prototype support itself is spatially variable. Normalization by $\log K$ places the maximum Jensen–Shannon divergence for $K$ states on a comparable 0–1 scale. The five-state curve remains the direct quantity used by `TP_HPM_full5`; the decomposition is reported to make its sources interpretable.

##### S5.3 Persistence integration and derived scales

Let $\delta = \frac{1}{2}\min\{\ell \in \mathcal{L} : \ell > 0\} = 0.0625$ mm and $u(\ell) = \log(1 + \ell/\delta)$. For $X \in \{5, +, \pm\}$,

$$TP - HPM_X = \frac{1}{u(\ell_{\max}) - u(0)} \sum_{m=1}^{|\mathcal{L}|-1} \frac{H_X(\ell_m) + H_X(\ell_{m+1})}{2} \left[u(\ell_{m+1}) - u(\ell_m)\right]. \tag{S25}$$

This is a normalized trapezoidal area under the heterogeneity-versus-log-scale curve. The characteristic radius $r_{50}$ is the first interpolated scale at which $H_X(\ell)$ falls to half its unsmoothed value. The fragmentation slope is the finite-difference slope between the first two scale points. Effective positive patch mass is $\sum_i q_i$, and reported physical area multiplies retained patch count by patch area at the acquisition resolution. Undefined statistics from degenerate slides were excluded only from the calculation requiring them; available fold-level estimates were averaged.

##### S5.4 Interpretation of the persistence curve

The unsmoothed value $H_X(0)$ quantifies fine-scale variability. As $\ell$ increases, local fluctuations are averaged and only broader spatial domains remain. A curve that falls rapidly indicates fragmented, patch-scale variation; a curve that remains high to 1–2 mm indicates larger coherent differences between tissue regions. Integrating on $u(\ell) = \log(1 + \ell/\delta)$ allocates resolution across small and large scales without allowing the longest linear interval to dominate solely because it is numerically wide.

Two slides can have the same persistence mass for different reasons. One may start at very high heterogeneity and decay rapidly, while another starts lower but persists. For this reason, the implementation also records $H(0)$, $r_{50}$, and the initial fragmentation slope. TP-HPM is a compact summary, not a sufficient description of the entire curve.

##### S5.5 Numerical checks

For every scale, probability entries were required to be finite and nonnegative, rows to sum to one, and Jensen–Shannon values to lie within their normalized theoretical ranges up to numerical tolerance. Connected tissue components remained invariant to smoothing scale. The numerical decomposition

$$\Delta_{\mathrm{decomp}}(\ell) = J_5(\ell) - J_\pm(\ell) - \bar{q}^{(\ell)} J_+(\ell) \tag{S26}$$

was required to remain close to zero. This QC residual detects inconsistent normalization, indexing, or state ordering. Coordinates and pixel size were validated before converting Gaussian widths to pixels, preserving the physical meaning of millimeter scales across scanners.

##### S5.6 Pseudocode summary

For each slide, the algorithm proceeds as follows: (1) validate the distance matrix, coordinates, and physical resolution; (2) rescale distances and calculate five-state probabilities; (3) split the spatial graph into connected tissue components; (4) smooth each probability channel at every physical scale using masked Gaussian normalization; (5) calculate five-state, positive-conditional, and positive-versus-negative Jensen–Shannon curves; (6) integrate each curve over the log-scale coordinate; and (7) export persistence masses, curve values, characteristic radii, fragmentation slopes, effective positive mass, and QC residuals. This deterministic sequence is implemented after prototype inference and has no survival outcome in its calculation.

### S6 Single-feature and prototype-family survival analyses

Multiple prototype features were tested under baseline, repulsion, reinitialization, and combined training. The largest contiguous region assigned to prototype 1 was associated with OS in BLOCAGE-01 across configurations (Table S4). A correlated six-feature family score was significant in BLOCAGE-01 and BCN (Table S5). These analyses showed that regional and correlated-family descriptors carried survival information and motivated the broader multifeature representation.

The closely aligned BLOCAGE-01 point estimates across training interventions demonstrate that the regional association was robust to prototype-geometry modifications. The broader family score extended the signal across two cohorts, while elastic-net integration allowed several complementary spatial families to contribute simultaneously.

Table S4: Largest prototype-1 region and OS in BLOCAGE-01, with age, sex, and KPS adjustment.

| Configuration | Unadjusted HR per SD | Unadjusted $P$ | Age/sex/KPS-adjusted $P$ |
|---|---|---|---|
| Baseline | 1.291 | 0.0008 | 0.0025 |
| Repulsion | 1.292 | 0.0008 | 0.0024 |
| Reinitialization | 1.298 | 0.0006 | 0.0024 |
| Combined | 1.288 | 0.0009 | 0.0040 |

Table S5: Correlated prototype-family score versus the single largest-region metric.

| Cohort | Slides (events) | Family-score HR ($P$) | Single-feature HR ($P$) |
|---|---|---|---|
| BLOCAGE-01 | 245 (101) | 1.305 (0.0119) | 1.290 (0.0008) |
| BCN | 41 (28) | 1.381 (0.0286) | 1.194 (0.2333) |
| LOC 2023 | 122 (68) | 0.982 (0.9099) | 0.924 (0.5888) |

The six-feature family score averaged correlated standardized metrics and reduced measurement noise across descriptors of the same latent structure. The multifeature elastic-net analysis below additionally permits coefficients of different signs and retains several spatial families.

### S7 Multifeature score derivation

#### S7.1 Correlation reduction and screening

Candidate metrics were grouped by absolute Spearman correlation, with distance $d_{jk} = 1 - |\rho_{jk}|$. Representative features were evaluated for correlation with KPS and for complementary survival information across cohorts. LOC 2023 and BLOCAGE-01 were pooled for coefficient estimation with cohort-stratified baseline hazards.

Absolute correlation groups features that move together or oppositely. If $\rho_{jk} \approx -1$, two metrics carry nearly the same rank information after sign reversal and were represented by one candidate. Family reduction lowered collinearity and the effective number of comparisons. KPS correlation screening reduced redundancy with performance status in the two development cohorts; the locked score was then applied unchanged to BCN.

##### S7.2 Standardization, stratified elastic net, and coefficients

For cohort $c$, feature $j$ was standardized by

$$Z_{ij} = \frac{X_{ij} - \bar{X}_{cj}}{s_{cj}}. \tag{S27}$$

The coefficient model used LOC 2023 and BLOCAGE-01 with cohort-stratified baseline hazards. Elastic-net regularization combines the sparsity of the lasso with the stabilization of ridge shrinkage [24–26]. With penalty $\lambda = 0.005$ and $l_1$ ratio $\alpha = 0.5$,

$$\widehat{\boldsymbol{\beta}} = \arg\min_{\boldsymbol{\beta}} \left\{ -\ell_{\text{stratified}}(\boldsymbol{\beta}) + \lambda \left[ \alpha \|\boldsymbol{\beta}\|_1 + \frac{1-\alpha}{2} \|\boldsymbol{\beta}\|_2^2 \right] \right\}, \tag{S28}$$

where

$$\ell_{\text{stratified}}(\boldsymbol{\beta}) = \sum_c \sum_{i \in \mathcal{D}_c} \left[ \boldsymbol{\beta}^\top \mathbf{Z}_i - \log \sum_{j \in \mathcal{R}_{ic}} \exp(\boldsymbol{\beta}^\top \mathbf{Z}_j) \right]. \tag{S29}$$

The ITH-C composite is $S_i = \widehat{\boldsymbol{\beta}}^\top \mathbf{Z}_i$. Table S6 reports the exact coefficients and mathematical interpretation of all 11 retained features.

Table S6: Exact ITH-C features, mathematical interpretation, and coefficients.

| Feature | Mathematical interpretation | Coefficient |
|---|---|---|
| `spatial_mixing_bin` | Indicator that unlike-prototype neighbor mixing $M$ exceeds its cohort median | −0.1166391914 |
| `tumor_confident_dominant_proto` | Numerical index $\arg\max_k f_k$ of the dominant positive prototype | −0.0413713435 |
| `TP_HPM_full5` | Normalized log-scale integral of five-state Jensen–Shannon heterogeneity $H_5(\ell)$ | −0.0000086834 |
| `proto0_minus_proto3` | Difference in positive-prototype fractions, $f_0 - f_3$ | +0.0000092580 |
| `tumor_confident_morans_I_bin_proto_3` | Moran's $I$ for the hard prototype-3 indicator field | +0.0150073987 |
| `morans_bin_tconf_mean` | Mean $4^{-1} \sum_{k=0}^{3} I_k$ across positive-prototype indicator fields | +0.0429577813 |
| `tumor_confident_mean_neighbours` | Mean spatial-graph degree among tumor-confident patches | +0.0513471582 |
| `tumor_confident_frac_proto_3` | Prototype-3 composition $f_3 = n_3 / \sum_k n_k$ | +0.1038651093 |
| `tumor_confident_frac_proto_2` | Prototype-2 composition $f_2 = n_2 / \sum_k n_k$ | +0.1338760491 |
| `tumor_confident_morans_I_bin_proto_1` | Moran's $I$ for the hard prototype-1 indicator field | +0.1498399939 |
| `q_bar_0` | Mean patch-level support assigned to positive prototype 0, $n^{-1} \sum_i p_{i0}$ | +0.1824026531 |

The near-zero raw coefficients for `TP_HPM_full5` and `proto0_minus_proto3` must be interpreted in the context of how those variables were stored and standardized. Coefficient magnitude alone should not be compared across unstandardized encodings.

For an individual slide, the standardized contribution of feature $j$ is $\widehat{\beta}_j Z_{ij}$. Summing these contributions recovers $S_i$. A positive coefficient means that, conditional on the other retained variables and the pooled stratified fit, larger standardized values increase fitted log hazard. This decomposition provides an inspectable contribution from every retained feature.

##### S7.3 Exact feature interpretations

1. `spatial_mixing_bin`: indicator that slide mixing $M$ exceeded its cohort median.
2. `tumor_confident_dominant_proto`: numerical index $\arg\max_k f_k$ of the most abundant positive prototype.
3. `TP_HPM_full5`: five-state persistence mass defined in the TP-HPM section.
4. `proto0_minus_proto3`: $f_0 - f_3$.
5. `tumor_confident_morans_I_bin_proto_3`: Moran's $I$ for the hard prototype-3 indicator field.
6. `morans_bin_tconf_mean`: mean of four hard-assignment Moran statistics.
7. `tumor_confident_mean_neighbours`: mean graph-neighbor count among retained tumor-confident patches.
8. `tumor_confident_frac_proto_3`: $f_3$.
9. `tumor_confident_frac_proto_2`: $f_2$.
10. `tumor_confident_morans_I_bin_proto_1`: Moran's $I$ for the hard prototype-1 indicator field.
11. `q_bar_0`: mean patch-level support assigned to positive prototype 0.

#### S8 Survival inference

For continuous standardized score $S_i^*$, adjusted Cox models were

$$h_i(t) = h_0(t) \exp\{\gamma S_i^* + \eta_1 \mathrm{age}_i + \eta_2 \mathrm{sex}_i + \eta_3 \mathrm{KPS}_i\}. \tag{S30}$$

A numerical ridge penalty of 0.001 was applied. The HR is $\exp(\widehat{\gamma})$ per within-cohort SD. Wald standard error $\mathrm{se}(\widehat{\gamma})$ gave

$$\mathrm{CI}_{95\%} = [\exp\{\widehat{\gamma} - 1.96\,\mathrm{se}(\widehat{\gamma})\}, \exp\{\widehat{\gamma} + 1.96\,\mathrm{se}(\widehat{\gamma})\}] \tag{S31}$$

[27]. The proportional-hazards assumption was evaluated with scaled Schoenfeld residuals. Kaplan–Meier estimates used

$$\widehat{S}(t) = \prod_{t_j \le t} \left(1 - \frac{d_j}{n_j}\right) \tag{S32}$$

[28]. Any score grouping in figures was used only for visualization and not as the primary inferential specification.

The partial likelihood used observed risk sets and did not require specification of $h_0(t)$. Primary cohort-specific adjusted models included age, sex, and continuous KPS; sensitivity models used age and sex or alternative age/KPS codings. A ridge penalty of 0.001 was added for numerical stability. The Wald interval uses the approximate normal distribution of the coefficient estimator.

The Kaplan–Meier median is the first time at which $\widehat{S}(t) \leq 0.5$. Square brackets denote the interquartile range (IQR), defined by times at which the curve crosses 0.75 and 0.25. If a curve does not cross a required probability before the final observation, that bound is reported as not reached.

Scaled Schoenfeld residuals were used to assess the proportional-hazards assumption in age-, sex-, and continuous-KPS-adjusted models. Correlation tests between the ITH-C residual process and transformed event time gave $P = 0.484$, $P = 0.115$, and $P = 0.718$ in LOC 2023, BLOCAGE-01, and BCN, respectively, providing no evidence that the ITH-C HR varied over follow-up.

##### S8.1 Development and external validation

LOC 2023 and BLOCAGE-01 contributed to feature-direction and penalty estimation and therefore yield apparent development estimates. The eleven-feature equation was locked before its application to BCN, which is the external international validation cohort. Cohort-specific estimates were displayed separately and were not statistically pooled, preserving the original development-versus-validation structure and avoiding a summary that conflates apparent and external effects. Top- and bottom-tertile contrasts were defined within cohort and fitted with the same age, sex, and continuous-KPS adjustment to express a clinician-interpretable separation while retaining the continuous score as the primary analysis. BCN clinical context was deliberately broad: 8 of 40 patients received autologous stem-cell transplantation, 8 were managed palliatively from the outset, and 4 received low-dose whole-brain radiotherapy, consistent with a real-life external cohort.

##### S8.2 Incremental value beyond MSKCC and KPS

MSKCC class was reconstructed according to the original definition: class I for age $\leq 50$ years, class II for age $> 50$ years and KPS $\geq 70$, and class III for age $> 50$ years and KPS $< 70$ [29]. The model portfolio compared ITH-C alone; MSKCC alone; MSKCC plus ITH-C; age plus sex; age plus sex plus ITH-C; age, sex, and continuous KPS; and the same clinical model plus ITH-C. Complete harmonized components of the IELSG score—serum lactate dehydrogenase, cerebrospinal-fluid protein, performance status, age, and deep-brain involvement—were not available across the development datasets, so an IELSG incremental-value analysis was not estimable.

For nested models, added value was tested by

$$\mathrm{LR} = 2\{\ell(\widehat{\theta}_{\mathrm{clinical+ITH-C}}) - \ell(\widehat{\theta}_{\mathrm{clinical}})\} \sim \chi^2_1. \tag{S33}$$

Discrimination gain was $\Delta C = C_{\mathrm{clinical+ITH-C}} - C_{\mathrm{clinical}}$, where $C$ is Harrell's concordance index [14]. Percentile 95% CIs used 1,000 patient-cluster bootstrap samples, preserving all slides from a patient in the same resample.

Continuous net reclassification improvement (NRI) and integrated discrimination improvement (IDI) were calculated at $t_0 = 24$ months with inverse-probability-of-censoring weights [30, 31]. With predicted event probability $\widehat{F}_m(t_0 \mid X_i)$ under model $m$, $\Delta_i = \widehat{F}_1 - \widehat{F}_0$, and weighted event/non-event expectations $E_w$, the continuous NRI was

$$\mathrm{NRI}(t_0) = E_w[\mathrm{sign}(\Delta_i) \mid T_i \le t_0] - E_w[\mathrm{sign}(\Delta_i) \mid T_i > t_0], \tag{S34}$$

and the IDI was the change in discrimination slope,

$$\mathrm{IDI}(t_0) = \{E_w(\widehat{F}_1 \mid T \le t_0) - E_w(\widehat{F}_1 \mid T > t_0)\} - \{E_w(\widehat{F}_0 \mid T \le t_0) - E_w(\widehat{F}_0 \mid T > t_0)\}. \tag{S35}$$

The censoring distribution was estimated within cohort. These reclassification measures are supportive because their bootstrap intervals were wider than those for $\Delta C$.

To assess whether the clinical-layer gain depended on refitting the same observations, fivefold patient-grouped cross-validation was repeated 50 times for MSKCC and MSKCC plus locked ITH-C. Test-fold C-indices were pooled within repetition. This sensitivity analysis cross-validates the downstream Cox coefficients; it does not re-estimate the upstream feature set or elastic-net penalty and is therefore not described as fully nested score development.

Table S7 presents the complete clinical comparison. ITH-C improved MSKCC discrimination from 0.671 to 0.717 in LOC 2023, from 0.560 to 0.593 in BLOCAGE-01, and from 0.588 to 0.706 in external BCN. The gains were supported by likelihood-ratio tests and remained positive in repeated downstream cross-validation. Adding ITH-C to age, sex, and continuous KPS increased $C$ from 0.748 to 0.754, 0.597 to 0.619, and 0.696 to 0.741, respectively. Figure S3 shows the consistency of HR estimates across alternative clinical-adjustment specifications without relying on the number of nominally significant fits.

Table S7: Survival association, proportional-hazards assessment, and clinical incremental value of ITH-C. **A)** Continuous and tertile contrasts.

| Cohort | Slides | Events | Age/sex/KPS HR per SD (95% CI); $P$ | PH test $P$ | Adjusted top vs bottom tertile HR (95% CI); $P$ |
|---|---|---|---|---|---|
| LOC 2023 | 122 | 68 | 1.29 (1.01–1.64); 0.0403 | 0.484 | 1.97 (1.01–3.84); 0.0467 |
| BLOCAGE-01 | 245 | 101 | 1.27 (1.07–1.51); 0.0055 | 0.115 | 1.78 (1.09–2.92); 0.0209 |
| BCN external | 41 | 28 | 2.13 (1.35–3.37); 0.0011 | 0.718 | 4.75 (1.60–14.11); 0.0050 |

Table S7: Survival association, proportional-hazards assessment, and clinical incremental value of ITH-C (continued). **B)** Harrell C-index model portfolio in the development and external-validation cohorts.

| Cohort | Model | Slides | Events | Harrell $C$ |
|---|---|---|---|---|
| LOC 2023 | ITH-C alone | 122 | 68 | 0.598 |
| | MSKCC | 122 | 68 | 0.671 |
| | MSKCC + ITH-C | 122 | 68 | 0.717 |
| | Age + sex | 122 | 68 | 0.717 |
| | Age + sex + ITH-C | 122 | 68 | 0.721 |
| | Age + sex + continuous KPS | 122 | 68 | 0.748 |
| | Age + sex + continuous KPS + ITH-C | 122 | 68 | 0.754 |
| BLOCAGE-01 | ITH-C alone | 245 | 101 | 0.546 |
| | MSKCC | 245 | 101 | 0.560 |
| | MSKCC + ITH-C | 245 | 101 | 0.593 |
| | Age + sex | 245 | 101 | 0.557 |
| | Age + sex + ITH-C | 245 | 101 | 0.582 |
| | Age + sex + continuous KPS | 245 | 101 | 0.597 |
| | Age + sex + continuous KPS + ITH-C | 245 | 101 | 0.619 |
| BCN external | ITH-C alone | 41 | 28 | 0.680 |
| | MSKCC | 41 | 28 | 0.588 |
| | MSKCC + ITH-C | 41 | 28 | 0.706 |
| | Age + sex | 41 | 28 | 0.661 |
| | Age + sex + ITH-C | 41 | 28 | 0.725 |
| | Age + sex + continuous KPS | 41 | 28 | 0.696 |
| | Age + sex + continuous KPS + ITH-C | 41 | 28 | 0.741 |

Table S7: Survival association, proportional-hazards assessment, and clinical incremental value of ITH-C (continued). **C)** Incremental-value statistics.

| Cohort | Reference model | $C_0$ | $C_1$ | $\Delta C$ (bootstrap 95% CI) | LR $P$ | NRI at 24 mo (95% CI) | IDI at 24 mo (95% CI) |
|---|---|---|---|---|---|---|---|
| LOC 2023 | MSKCC | 0.671 | 0.717 | 0.046 (0.006–0.095) | 0.027 | 0.318 (−0.104–0.917) | 0.038 (−0.000–0.134) |
| | MSKCC + sex | 0.688 | 0.722 | 0.034 (−0.001–0.087) | 0.033 | 0.244 (−0.121–0.879) | 0.035 (0.000–0.126) |
| | Age + sex + continuous KPS | 0.748 | 0.754 | 0.005 (−0.007–0.043) | 0.041 | 0.626 (−0.063–1.003) | 0.030 (−0.000–0.093) |
| BLOCAGE-01 | MSKCC | 0.560 | 0.593 | 0.033 (0.002–0.085) | 0.008 | 0.210 (−0.025–0.535) | 0.020 (−0.000–0.061) |
| | MSKCC + sex | 0.563 | 0.597 | 0.033 (−0.002–0.083) | 0.008 | 0.210 (−0.015–0.585) | 0.020 (−0.000–0.071) |
| | Age + sex + continuous KPS | 0.597 | 0.619 | 0.022 (−0.003–0.071) | 0.008 | 0.212 (−0.035–0.561) | 0.019 (−0.001–0.064) |
| BCN external | MSKCC | 0.588 | 0.706 | 0.118 (0.017–0.205) | 0.003 | 0.370 (−0.348–1.153) | 0.145 (0.007–0.318) |
| | MSKCC + sex | 0.599 | 0.702 | 0.102 (0.007–0.229) | 0.004 | 0.370 (−0.446–1.214) | 0.141 (0.006–0.312) |
| | Age + sex + continuous KPS | 0.696 | 0.741 | 0.045 (−0.013–0.156) | 0.001 | 0.459 (−0.208–1.239) | 0.158 (0.015–0.298) |

$C_0$ and $C_1$ denote the clinical reference model without and with ITH-C, respectively. Bootstrap intervals used 1,000 patient-cluster resamples. NRI, continuous net reclassification improvement; IDI, integrated discrimination improvement; LR, likelihood-ratio test.

Table S7: Survival association, proportional-hazards assessment, and clinical incremental value of ITH-C (continued). **D)** Repeated patient-grouped downstream cross-validation of MSKCC models.

| Cohort | MSKCC mean $C$ | MSKCC + ITH-C mean $C$ | Mean $\Delta C$ (2.5th–97.5th percentile) | Repetitions |
|---|---|---|---|---|
| LOC 2023 | 0.670 | 0.710 | 0.040 (0.023–0.059) | 50 |
| BLOCAGE-01 | 0.559 | 0.584 | 0.026 (0.009–0.039) | 50 |
| BCN external | 0.554 | 0.676 | 0.122 (0.027–0.246) | 50 |

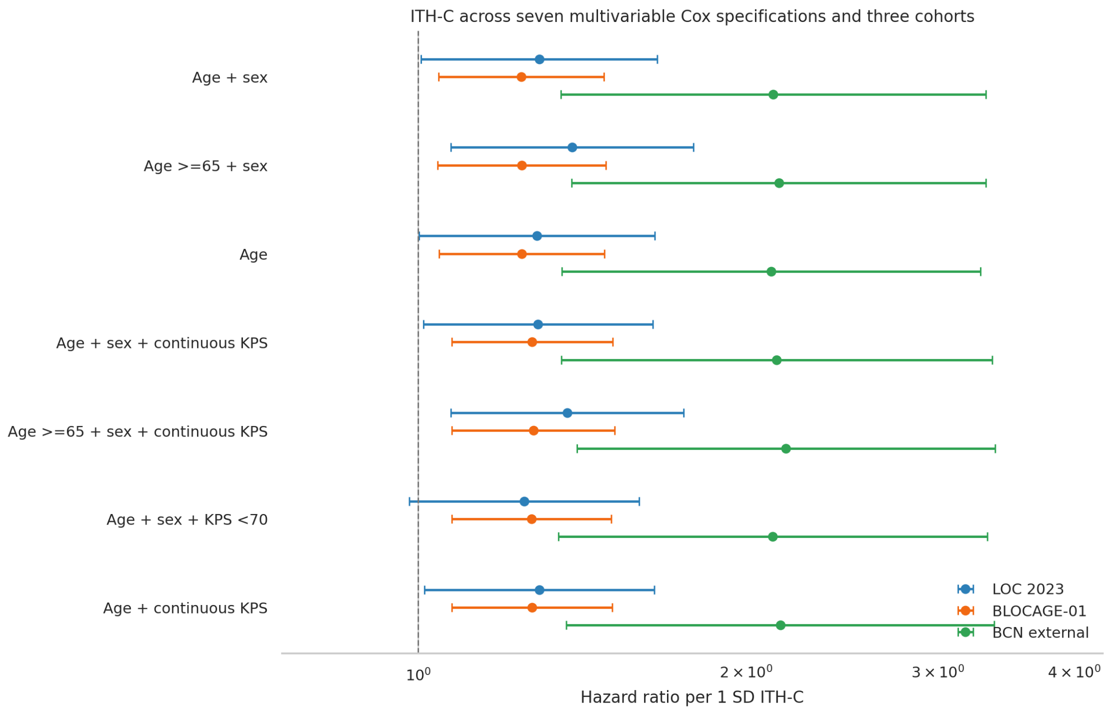

**Fig. S3. ITH-C hazard ratios across seven Cox-model specifications and three cohorts.** Hazard ratios are reported per one standard deviation increase in ITH-C on a logarithmic scale. Blue, orange, and green denote LOC 2023, BLOCAGE-01, and BCN, respectively. Continuous and dichotomized KPS specifications were evaluated in all three cohorts.

### S9 Eight-biopsy spatial transcriptomic analysis

#### S9.1 Datasets, alignment, and expression preprocessing

Eight treatment-naive PCNSL Visium sections were analyzed. P1, P2, P3, and P7 were obtained from GSE203552 [32]; Hot, Cold, immune-microenvironment-excluded (IME), and immune-microenvironment-suppressed (IMS) were obtained from GSE230207 [33]. The combined dataset contained 22,532 tissue spots (Table S8). Count-matrix barcodes were intersected with the tissue-position table, and H&E images were checked against spot-array orientation. Spots outside tissue or failing expression quality control were removed. Counts were library normalized and log transformed for expression summaries and marker scoring; Starfysh received the count representation required by its generative model.

Table S8: Public PCNSL Visium sections included in the spatial analysis.

| Section | GEO sample | Study accession | Tissue spots | Detected genes |
|---|---|---|---|---|
| P1 | GSM6176208 | GSE203552 | 1,083 | 17,943 |
| P2 | GSM6176213 | GSE203552 | 917 | 17,943 |
| P3 | GSM6176214 | GSE203552 | 1,512 | 17,943 |
| P7 | GSM6176215 | GSE203552 | 1,090 | 17,943 |
| Hot | GSM7192449 | GSE230207 | 4,517 | 17,922 |
| Cold | GSM7192450 | GSE230207 | 4,395 | 17,922 |
| IME | GSM7192451 | GSE230207 | 4,349 | 17,922 |
| IMS | GSM7192453 | GSE230207 | 4,669 | 17,922 |

##### S9.2 Starfysh CS1–CS4 mapping

Starfysh is a histology-aware deep generative model that represents each Visium spot as a mixture of marker-anchored expression programs [34]. After selection of 2,000 highly variable genes, the supplied 100-gene CS1–CS4 lists from the PCNSL consensus study were intersected with expressed genes and supplied as anchors [16]. Table S9 summarizes the four source lists and their leading genes. For normalized spot $s$, posterior program support was represented by $\boldsymbol{\pi}_s = (\pi_{s1}, \ldots, \pi_{s4})$, $\pi_{sk} \geq 0$, and $\sum_k \pi_{sk} = 1$. The model reconstructs gene counts through program-specific latent expression while regularizing the mixture with marker and spatial-histology information. Three random initializations were run per section using seeds 20260818–20260820; the fit with the lowest final reconstruction objective was retained. Posterior means were returned to native spot coordinates. The full eight-section H&E, CS1–CS4, and local ITH-C atlas is Figure 5 in the main manuscript.

Table S9: Source CS1–CS4 anchor lists supplied to the spatial workflow.

| Program | Source genes | Representative genes in supplied rank order |
|---|---|---|
| CS1 | 100 | *IGHV3-64D, PPP1R2C, RFPL4AP1, IGLV7-43, HOXD11, SSX5, IGHV4-80, CAMTA1-AS1, CT45A5, KRT8P5, IGHV3-60, RPL36AP40* |
| CS2 | 100 | *RNU6-329P, OR10T1P, RNU6-535P, RN7SL843P, MED14P1, RNA5SP349, RPS12P31, LINC02140, ATP6V0E1P1, HTN3, USP9YP3, MTND3P9* |
| CS3 | 100 | *IGKV1-17, YRDCP1, IGHV3-73, TRBV12-5, LINC01228, TBL1XR1-AS1, VTRNA1-2, PRR15L, HMGB1P17, IGKV1-6, RN7SKP211, FBP1* |
| CS4 | 100 | *ACOD1, TRAV12-2, CXCR2P1, TRAJ22, KLRC1, TRAV12-1, TRBV4-1, GNLY, LINC01871, GBP5, FCRL6, TRAV5* |

Counts refer to the source RBraLymP-derived lists before intersection with the 2,000 highly variable genes and the genes measured in each Visium study.

CS1–CS4 were derived from bulk RNA and combine malignant B-cell and microenvironmental biology. Consequently, a local Starfysh proportion denotes support for a tissue-ecosystem program, not a purified malignant-cell subtype. A fifth low-CS state, $p_{s5} = 1 - \max_k \pi_{sk}$ after within-section calibration, represented limited positive CS support and completed the normalized five-state field $\mathbf{p}_s = (p_{s1}, \ldots, p_{s5})$ used below.

##### S9.3 Spot-level domain-adapted ITH-C projection

Spot coordinates were converted to millimeters. The undirected spatial graph connected at most six nearest neighbors within 0.13 mm. Let $A = (a_{ij})$ be its symmetric adjacency matrix and $\mathcal{N}_i$ the neighbors of spot $i$. Eleven local terms were reconstructed using the same coefficients as the WSI ITH-C model: binarized local mixing, dominant state, local five-state persistence mass, prototype-0 minus prototype-3 support, local Moran contributions for prototype 3 and their mean, mean positive-state degree, fractions of prototypes 3 and 2, the local Moran contribution for prototype 1, and $q_{i0} = p_{i0}$. For each term $x_{ij}$, within-section standardization was

$$z_{ij} = \frac{x_{ij} - \bar{x}_{\cdot j}}{s_j}. \tag{S36}$$

The unsmoothed local projection was

$$\begin{aligned} R_i = & -0.116639 z_{i,\text{mix}} - 0.041371 z_{i,\text{dominant}} - 0.000008683 z_{i,\text{HPM5}} + 0.000009258 z_{i,p_0-p_3} \\ & + 0.015007 z_{i,I_3} + 0.042958 z_{i,\bar{I}} + 0.051347 z_{i,d^+} + 0.103865 z_{i,f_3} \\ & + 0.133876 z_{i,f_2} + 0.149840 z_{i,I_1} + 0.182403 z_{i,q_0}. \end{aligned} \tag{S37}$$

Here, local mixing is $|\mathcal{N}_i|^{-1} \sum_{j \in \mathcal{N}_i} \mathbf{1}(c_i \neq c_j)$; $d_i^+ = \sum_j a_{ij} \mathbf{1}(c_j < 5)$; and $I_{ik}$ is the local contribution of the indicator $\mathbf{1}(c_i = k)$ to global Moran autocorrelation. Local HPM5 integrated normalized Jensen–Shannon divergence across $\ell \in \{0, 0.125, 0.25, 0.5, 1, 2\}$ mm using the same log-scale trapezoidal integration described for WSI TP-HPM.

$R_i$ was standardized within section, smoothed with a 0.125-mm Gaussian kernel separately inside each connected tissue component, and standardized again:

$$\text{ITHC}_i^{\text{Visium}} = \mathcal{Z}\left[\frac{\sum_{j \in C(i)} K_{0.125}(d_{ij})\, \mathcal{Z}(R_j)}{\sum_{j \in C(i)} K_{0.125}(d_{ij})}\right], \qquad K_h(d) = \exp\left(-\frac{d^2}{2h^2}\right), \tag{S38}$$

where $C(i)$ is the connected tissue component containing spot $i$. This exact reconstruction reproduced all 22,532 cached local scores with maximum absolute error $2.4 \times 10^{-13}$. The spatial score is denoted $\text{ITHC}^{\text{Visium}}$ to distinguish it from the clinical WSI ITH-C.

##### S9.4 Immune-compartment enrichment and cell2location analyses

Nine readable compartments were evaluated: tumor/B-cell program, $CD4^+$ helper T cells, $CD8^+$ cytotoxic T cells, regulatory T cells, natural-killer cells, resident microglia-like cells, monocyte-derived macrophages, conventional dendritic cells, and plasma cells. Marker genes were assigned to 24 expression bins, and every marker set was matched to control genes with similar mean expression. For compartment $c$ at spot $i$, the control-subtracted expression and detection contrasts were

$$E_{ic} = \overline{Z(Y_{ig})}_{g \in M_c} - \overline{Z(Y_{ig})}_{g \in C_c}, \qquad D_{ic} = \overline{\mathbf{1}(X_{ig} > 0)}_{g \in M_c} - \overline{\mathbf{1}(X_{ig} > 0)}_{g \in C_c}. \tag{S39}$$

The consensus enrichment was $U_{ic} = 0.75 E_{ic} + 0.25 D_{ic}$ and relative spot weights used a softmax temperature of 1.25. These quantities are relative enrichments, not absolute cell fractions. Spatial maps are shown in Figure S4.

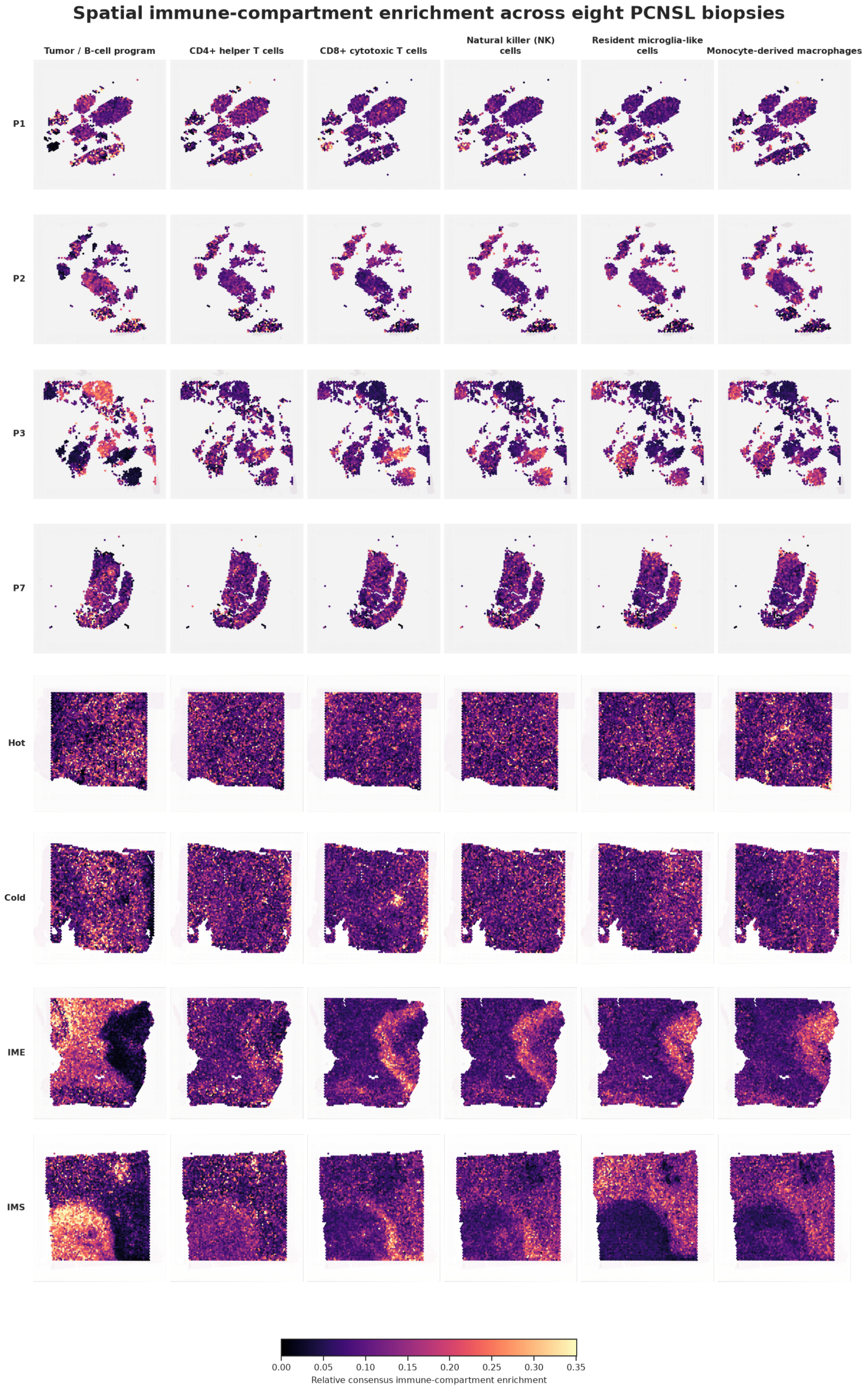

**Fig. S4. Relative immune-compartment enrichment across eight PCNSL Visium biopsies.** Rows denote sections and columns denote the tumor/B-cell and principal immune programs. Scores combine expression-matched marker-control subtraction and marker-detection enrichment and are visualized on one common relative scale.

Complementary cell2location deconvolution used the matched GSE203552 single-cell RNA-sequencing atlas [32, 35]. After quality control, 73,896 cells were represented by an NB2 reference-regression model across 9,036 retained genes and 11 cell types. The reference model was trained for 150 epochs. Spatial mapping used the 2,000 most cell-type-informative shared genes, $N_{\text{cells/location}} = 20$, detection $\alpha = 200$, batch size 2,048, 700 epochs, and 500 posterior samples. The fifth-percentile posterior abundance (q05) provided a conservative per-spot abundance estimate. Table S10 reports the exact reference taxonomy and the marker rules used to refine T-cell and myeloid annotations.

Table S10: Cell2location reference taxonomy and annotation rules.

| Reference cell type | Assignment rule | Marker genes or source annotation |
|---|---|---|
| Malignant B cell | Original study annotation | GSE203552 `mBc` |
| Normal B cell | Original study annotation | GSE203552 `nmBc` |
| CD4 helper T cell | Highest T-cell subtype score | *CD3D, CD3E, IL7R, CD4, LTB* |
| CD8 cytotoxic T cell | Highest T-cell subtype score | *CD8A, CD8B, GZMK, NKG7, CCL5* |
| Regulatory T cell | Highest T-cell subtype score | *FOXP3, IL2RA, CTLA4, TIGIT* |
| Natural-killer cell | Highest T-cell subtype score | *GNLY, KLRD1, PRF1, NCAM1, KLRF1* |
| Proliferating T cell | Highest T-cell subtype score | *MKI67, TOP2A* |
| Microglia | Microglia score > MDM score | *CX3CR1, P2RY12, TMEM119, SALL1, GPR34* |
| Monocyte-derived macrophage | MDM score ≥ microglia score | *CD14, FCGR3A, S100A8, S100A9, VCAN, LST1* |
| Dendritic cell | Original study annotation | GSE203552 `mDC1` |
| Oligodendrocyte | Original study annotation | GSE203552 `oligo` |

Five tumor–immune-interface definitions were derived only from cell2location and coordinates: malignant/immune rank balance, co-abundance, balanced co-abundance, local malignant-fraction gradient, and equidistance from high-confidence malignant and immune cores. Their independence from the ITH-C equation enabled a direct spatial cross-check. The q05 abundances and marker assignments were further evaluated through driver-association and leave-one-section-out sensitivity analyses.

#### S9.5 RNA-inferred CNV, directional burden, and aneuploidy

Gene coordinates were assigned from GENCODE v50 (GRCh38) [36]. infercnvpy 0.6.1 [37] averaged expression-ordered genes using a 100-gene window and 10-gene step, excluded chromosomes X and Y, clipped log-fold change at 3, and applied a dynamic threshold of 1.5. With no unequivocal normal spatial reference shared by all sections, the reference was the within-section mean. For bin $b$ and spot $i$, the smoothed relative signal was denoted $v_{ib}$. Directional and total burdens were

$$G_i = B^{-1} \sum_b \max(v_{ib}, 0), \quad L_i = B^{-1} \sum_b \max(-v_{ib}, 0), \quad T_i = G_i + L_i. \tag{S40}$$

The altered-bin fraction was $B^{-1} \sum_b \mathbf{1}(|v_{ib}| > t)$, where $t$ was the infercnvpy dynamic threshold. Chromosome-level means were $\bar{v}_{ic} = |\mathcal{B}_c|^{-1} \sum_{b \in \mathcal{B}_c} v_{ib}$, and the continuous aneuploidy index was $A_i = 22^{-1} \sum_{c=1}^{22} |\bar{v}_{ic}|$. Between 1,511 and 1,513 autosomal bins were retained per section. Gain, loss, aneuploidy, and local ITH-C fields are shown in Figure S5. These are relative RNA-inferred patterns and not DNA ploidy or DNA-validated copy-number calls.

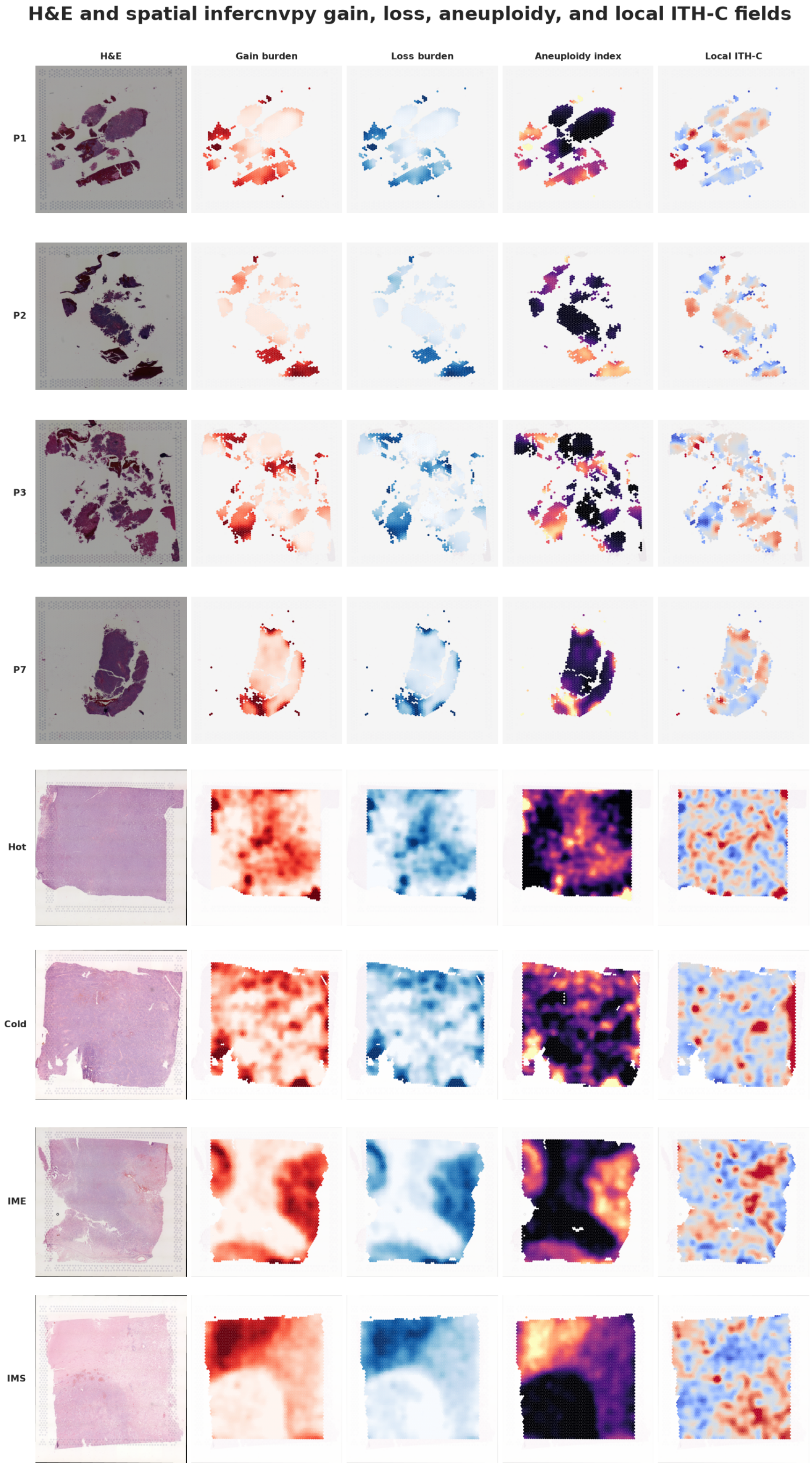

**Fig. S5. Spatial RNA-inferred chromosomal alteration fields.** Each row shows H&E, relative gain burden, relative loss burden, the continuous aneuploidy index, and local $\mathrm{ITHC}^{\mathrm{Visium}}$. infercnvpy values are centered within section and identify regional expression imbalance rather than absolute DNA copy number.

##### S9.6 T-cell, macrophage, and checkpoint-state signatures

Nine curated immune-state signatures were scored with Scanpy 1.12.3 `score_genes`. Each intended list was intersected with measured genes and required at least three available genes. With normalized log-expression $Y_{ig}$, the score for signature $s$ at spot $i$ was

$$S_{is} = \frac{1}{|G_s|} \sum_{g \in G_s} Y_{ig} - \frac{1}{|C_s|} \sum_{h \in C_s} Y_{ih}, \tag{S41}$$

where $G_s$ is the available signature set and $C_s$ is the expression-matched control set selected with the Scanpy defaults of 25 expression bins and control size 50. Table S11 provides the complete intended gene definitions. These state scores are relative expression contrasts and not cell fractions.

Table S11: Curated immune-state gene signatures used in the Visium analysis.

| Signature | Interpretation | Intended genes before intersection with measured genes |
|---|---|---|
| T effector | Cytotoxic effector function | *GZMB, PRF1, IFNG, NKG7, GNLY, GZMA, GZMK, FGFBP2* |
| T exhaustion | T-cell dysfunction/exhaustion | *PDCD1, HAVCR2, LAG3, TIGIT, TOX, CTLA4, HLA-DRA* |
| T activation | Recent activation and costimulation | *CD69, ICOS, IL2RA, TNFRSF4, TNFRSF9* |
| T proliferation | Cell-cycle activity | *MKI67, TOP2A, BIRC5, PCNA* |
| T naive/memory | Naive or central-memory state | *CCR7, SELL, TCF7* |
| T terminal effector | Terminal cytotoxic differentiation | *GZMB, PRF1, FGFBP2, FCGR3A, SPON2* |
| M1 proinflammatory | Proinflammatory myeloid program | *IL1B, TNF, IL6, CXCL9, CXCL10, NOS2, SOCS3, CCL5* |
| M2/TAM | Tumor-associated macrophage program | *CD163, MRC1, CCL22, TREM2, SPP1, GPNMB, CD9, CLEC7A* |
| Checkpoint ligands | Immune-checkpoint ligand program | *CD274, PDCD1LG2, CD47, LGALS9* |

Dominant CS groups were assigned from the maximum Starfysh posterior. ITH-C tiers were defined separately within each section using tertiles, yielding 7,511 low, 7,510 middle, and 7,511 high spots. Violin plots display pooled spot distributions, and omnibus differences used the Kruskal–Wallis test. Medians, interquartile ranges, group sizes, statistics, and exact numerical values are reported in Tables S12 and S13. Section-aware meta-analysis and paired pseudobulk contrasts provided the principal biological replication framework.

##### S9.7 Association models and section-aware inference

Within each section, ordinary Spearman correlations and partial Spearman correlations were calculated after rank residualization for log library size and RNA-inferred total CNV burden. Spot-level section estimates were Fisher transformed, $z_s = \operatorname{atanh}(r_s)$, and combined using inverse variance $w_s = n_s - 3$:

$$r_{\text{meta}} = \tanh\left(\frac{\sum_s w_s z_s}{\sum_s w_s}\right). \tag{S42}$$

Benjamini–Hochberg correction was applied within each analysis family. Because spots within a section are spatially dependent, complementary section-level correlations, random-effects summaries, and exact sign-flip tests across the eight sections were also calculated. High-versus-low ITH-C pseudobulks used section-specific tertiles. Counts were summed within tier, converted to counts per million, and contrasted within section. Direction consistency was emphasized; no gene passed sign-test FDR correction with only eight paired sections.

The principal association results are summarized in Figure S6. Partial meta-correlations of local ITH-C were positive for $CD8^+$ cytotoxic T cells ($r = 0.138$), natural-killer cells ($r = 0.134$), resident microglia-like cells ($r = 0.036$), and monocyte-derived macrophages ($r = 0.064$), and negative for the tumor/B-cell program ($r = -0.152$) and plasma cells ($r = -0.139$). High-ITH-C pseudobulks showed concordant increases in cytotoxic and interferon/myeloid genes, including *CD8A*, *IFNG*, *GZMA*, *CXCL9*, *LAG3*, *C1QA*, and *C1QB*, across all eight sections. RNA-inferred gain, loss, and aneuploidy had small negative spot-level correlations with local ITH-C ($r = -0.041$, $-0.071$, and $-0.043$), while section-level aneuploidy was unrelated to mean ITH-C ($\rho = -0.12$, $P = 0.779$).

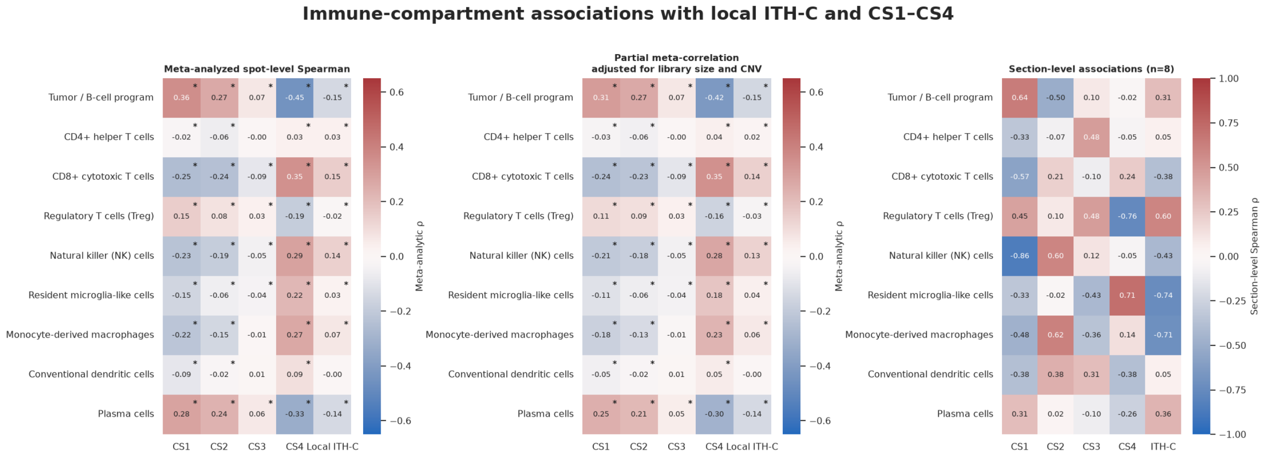

**(a)** Immune associations.

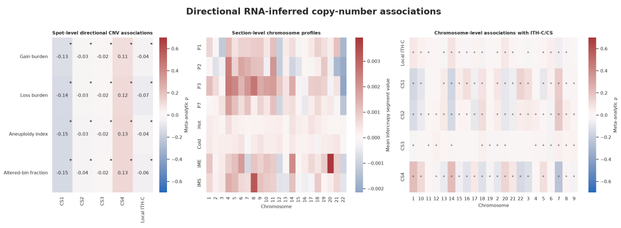

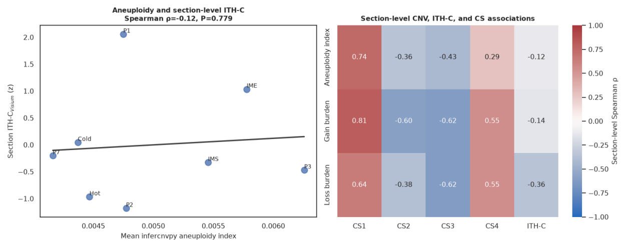

**(b)** Directional CNV associations.

**(c)** Aneuploidy associations.

**Fig. S6. Immune, directional copy-number, and aneuploidy associations with CS1–CS4 and local ITH-C. a)** Ordinary and partial Fisher-$z$ meta-correlations and section-level correlations. **b)** Spot-level gain and loss meta-correlations and chromosome-level profiles. **c)** Section-level aneuploidy, ITH-C, and CS relationships. Asterisks indicate false-discovery-rate-adjusted $P < 0.05$.

Table S12: T-cell signature scores by dominant CS1–CS4 program.

| Signature | CS1 ($n = 6{,}344$) | CS2 ($n = 5{,}880$) | CS3 ($n = 4{,}626$) | CS4 ($n = 5{,}682$) | Kruskal $H$ | $P$ |
|---|---|---|---|---|---|---|
| T effector | −0.209 [0.150] | 0.229 [0.259] | −0.079 [0.343] | 0.241 [0.492] | 8,390.67 | $< 10^{-300}$ |
| T exhaustion | 0.066 [0.207] | 0.207 [0.182] | 0.061 [0.241] | 0.263 [0.220] | 5,679.51 | $< 10^{-300}$ |
| T activation | −0.048 [0.154] | −0.079 [0.159] | −0.080 [0.153] | 0.011 [0.190] | 1,401.90 | $1.14 \times 10^{-303}$ |
| T proliferation | 0.189 [0.332] | −0.062 [0.292] | −0.044 [0.332] | −0.165 [0.448] | 4,296.27 | $< 10^{-300}$ |
| T naive/memory | −0.009 [0.287] | 0.141 [0.325] | −0.080 [0.318] | 0.010 [0.267] | 1,306.01 | $7.30 \times 10^{-283}$ |
| T terminal effector | −0.221 [0.237] | 0.134 [0.241] | 0.049 [0.440] | 0.417 [0.460] | 8,065.15 | $< 10^{-300}$ |

Values are median [interquartile range] Scanpy control-adjusted expression scores. The omnibus Kruskal–Wallis test uses pooled spots; section-aware analyses are described in the text.

Table S13: T-cell signature scores by within-section ITH-C tier.

| Signature | Low ($n = 7{,}511$) | Middle ($n = 7{,}510$) | High ($n = 7{,}511$) | Kruskal $H$ | $P$ |
|---|---|---|---|---|---|
| T effector | −0.102 [0.420] | −0.005 [0.442] | 0.173 [0.520] | 1,535.65 | $< 10^{-300}$ |
| T exhaustion | 0.083 [0.257] | 0.155 [0.215] | 0.219 [0.236] | 2,337.07 | $< 10^{-300}$ |
| T activation | −0.069 [0.156] | −0.054 [0.164] | −0.028 [0.189] | 344.35 | $1.68 \times 10^{-75}$ |
| T proliferation | 0.042 [0.336] | 0.015 [0.354] | −0.121 [0.451] | 965.70 | $2.00 \times 10^{-210}$ |
| T naive/memory | 0.014 [0.347] | 0.060 [0.340] | −0.018 [0.322] | 299.35 | $9.92 \times 10^{-66}$ |
| T terminal effector | −0.035 [0.413] | 0.032 [0.417] | 0.268 [0.579] | 1,953.52 | $< 10^{-300}$ |

Values are median [interquartile range] Scanpy control-adjusted expression scores. Tiers are section-specific tertiles, preserving equal representation of each biopsy across the pooled comparison.

#### S9.8 Moran and fixed-lattice Ripley-type spatial analyses

Permutation Moran's $I$ used the section-specific Visium graph. For centered feature vector $\mathbf{x}$ and total edge weight $W = \sum_{ij} a_{ij}$,

$$I = \frac{n}{W} \frac{\sum_{ij} a_{ij}(x_i - \bar{x})(x_j - \bar{x})}{\sum_i (x_i - \bar{x})^2}. \tag{S43}$$

Two-sided permutation $P$ values used 199 label permutations. Moran analyses included local ITH-C, CS1–CS4, gain, loss, aneuploidy, and selected immune compartments. Local ITH-C was significant in every section ($I = 0.559$–$0.939$), as were gain, loss, and aneuploidy fields (Figure S7).

For fixed-lattice Ripley-type analysis, feature values were converted to a high-state indicator $y_i = \mathbf{1}(x_i \geq Q_{0.75})$. At radius $r$, the observed number of high–high unordered spot pairs was

$$Q_{\text{obs}}(r) = \sum_{i<j} \mathbf{1}(d_{ij} \leq r) y_i y_j. \tag{S44}$$

The same number of high spots was reassigned to the fixed lattice in each of 199 permutations. Enrichment was

$$E(r) = \frac{Q_{\text{obs}}(r)}{199^{-1} \sum_{b=1}^{199} Q_b(r)} - 1, \tag{S45}$$

evaluated at 0.125, 0.25, 0.5, 1, and 2 mm. The one-sided permutation $P$ value was $(1 + \sum_b \mathbf{1}[Q_b \geq Q_{\text{obs}}])/200$, followed by Benjamini–Hochberg correction. This fixed-lattice formulation tests aggregation of high values without changing tissue geometry and is related to second-order point-pattern analysis [38].

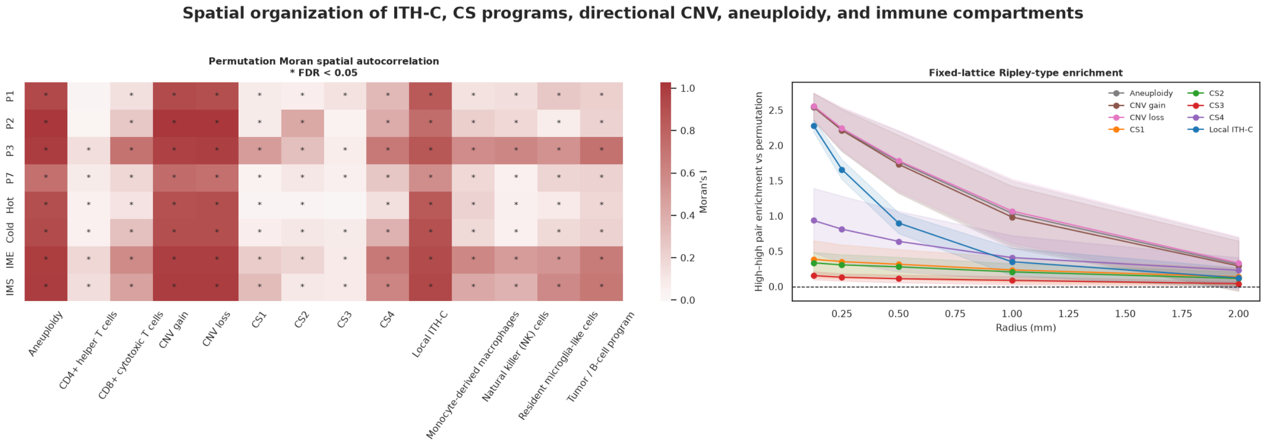

**Fig. S7. Spatial autocorrelation and multiscale high–high clustering.** Left, permutation Moran's $I$ by section for local ITH-C, CS programs, RNA-inferred chromosomal alteration fields, and immune compartments. Right, across-section mean fixed-lattice Ripley-type enrichment with 95% standard-error bands.

The section-resolved curves are presented in Figure 6a of the main manuscript, avoiding duplication with the supplementary summary in Figure S7. At 0.125 mm, median enrichment was 2.283 for local ITH-C, 2.539 for gain, 2.608 for loss, and 2.586 for aneuploidy; all were significant in eight of eight sections. CS-specific median enrichment was 0.158, 0.352, 0.193, and 0.862 for CS1–CS4, respectively. Section-level estimates at every radius were used for the plotted curves and multiplicity-adjusted inference.

#### S9.9 T-cell state sensitivity analyses

The CS- and ITH-C-stratified violin plots are presented in Figure 6b–c of the main manuscript and quantified in Tables S12 and S13. CS4-dominant spots had the highest median effector, exhaustion, activation, and terminal-effector scores; CS1 had the highest proliferation score. Across ITH-C tiers, effector, exhaustion, activation, and terminal-effector medians rose progressively, whereas proliferation declined. The joint effector–exhaustion distribution and spatial maps retained for the Supplementary methods are shown in Figure S8. This allocation provides complementary spatial detail without repeating the main panels.

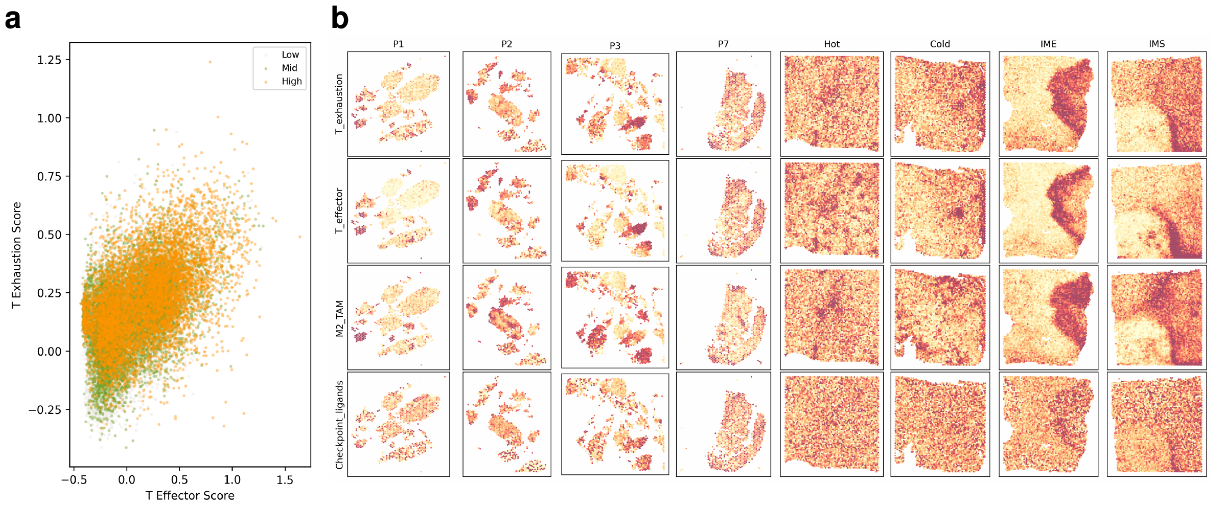

**Fig. S8. Complementary T-cell and immune-state sensitivity analyses. a)** Joint T-cell effector and exhaustion scores colored by within-section ITH-C tier. **b)** Spatial maps of T-cell exhaustion, T-cell effector, M2/TAM, and checkpoint-ligand scores across all eight biopsies. Scores are within-dataset Scanpy control-adjusted expression contrasts.

### S10 Software, code, and reproducibility checklist

- Python version: 3.10.0 (conda environment `clam_latest`).
- Core libraries: PyTorch, torchvision, OpenSlide, CLAM, UNI, NumPy, pandas, SciPy, scikit-learn, lifelines, pycox, matplotlib, and seaborn.
- Exact package versions: PyTorch 2.9.1, torchvision 0.24.1, OpenSlide 4.0.0 (via openslide-python 1.4.3), NumPy 2.2.6, pandas 2.3.3, SciPy 1.15.3, scikit-learn 1.7.2, lifelines 0.30.0, pycox 0.3.0, matplotlib 3.10.8, seaborn 0.13.2.
- Spatial-CNV environment: Python 3.12.13, Starfysh 1.2.0, Scanpy 1.12.3, infercnvpy 0.6.1, PyTorch 2.13.0, pandas 2.3.3, matplotlib 3.10.9, and seaborn 0.13.2; random seed 20260818.
- Spatial association environment: Python 3.11.14, NumPy 2.4.6, pandas 2.3.3, SciPy 1.17.1, scikit-learn 1.8.0, matplotlib 3.10.9, and seaborn 0.13.2; random seed 20260821.
- Cell2location configuration: NB2 reference regression over 9,036 genes and 11 cell types; 150 reference epochs; 2,000 spatial genes; 700 spatial epochs; batch size 2,048; 500 posterior samples; q05 abundances; random seed 20260818.
- Public analysis code and trained weights: `https://github.com/lucas-rdlr/PCNSL-ITHC`.
- Immutable archive: code – `https://doi.org/10.5281/zenodo.22180418`; trained model checkpoints – `https://doi.org/10.5281/zenodo.21886774`.

#### S11 Methodologic interpretation and translational scope

The full pipeline converts routine WSIs into an outcome-associated spatial phenotype through four connected levels: fixed patch representation, weakly supervised outcome and consensus-program learning, explicit prototype probability fields, and an elastic-net combination of mathematically defined heterogeneity features. This design preserves the representational power of deep learning while exposing the composition, topology, autocorrelation, and persistence variables that contribute to ITH-C.

Technical portability was demonstrated across 40× Hamamatsu NDPI and 3DHISTECH MRXS acquisitions and across tissue contributed by more than 10 French reference centers and one Spanish center. Physical-coordinate conversion ensured that neighborhoods and Gaussian smoothing scales represented the same distances in millimeters despite differences in native pixel size. The consistent ITH-C association across these settings supports extension to additional scanners and institutions through fixed preprocessing, feature-schema validation, and reference normalization.

The microenvironment is an integral component of the learned phenotype. Bulk consensus signatures incorporate immune and stromal biology, and spatial immune organization can itself carry prognostic information [39]. The five-state field captures both positive-prototype composition and the negative/background alternative; its Jensen–Shannon decomposition further separates variation among CS-associated states from variation in total positive-prototype support. This integrated view is particularly well suited to PCNSL, in which malignant B cells, reactive brain, vessels, and recruited immune populations form tightly coupled spatial ecosystems.

The framework is directly extensible to other tumors. Disease-relevant prototype states can be combined with the same global diversity, local neighborhood, autocorrelation, connected-region, and multiscale persistence operators. Because the final Cox score remains decomposable into signed standardized contributions, applications to brain tumors and pan-cancer cohorts can retain interpretability while benefiting from increasingly capable pathology foundation models.
